# *In vivo* intercellular CRISPR screens using viral proximity barcoding reveal regulators of tumor-immune interactions

**DOI:** 10.64898/2026.09.16.752238

**Authors:** Peter P. Du, Seth T. Kohno, Mengchen Wang, Michael Papanicolaou, Alun Vaughan-Jackson, Leighton H. Daigh, Qiangwei Peng, Kaitlyn Spees, Aaron K. McCormick, Markus I. Diehl, Lacramioara Bintu, Xiaojie Qiu, Ansuman T. Satpathy, Michael C. Bassik

**Affiliations:** Cancer Biology Program, Stanford University School of Medicine, Stanford, CA, USA; Department of Genetics, Stanford University School of Medicine, Stanford, CA, USA; Department of Pathology, Stanford University School of Medicine, Stanford, CA, USA; Department of Computer Science, Stanford University, Stanford, CA, USA; Chan Zuckerberg Biohub, San Francisco, CA, USA; Department of Bioengineering, Stanford University, Stanford, CA, USA; Immunology Program, Stanford University School of Medicine, Stanford, CA, USA

## Abstract

Cell-cell interactions shape tumor growth, immune evasion, and therapeutic response, but systematically dissecting their genetic regulators *in vivo* remains challenging. Existing cell-cell interaction tracing technologies rely largely on protein labels or enzymatic reactions, limiting their information content and scalability. To overcome these limitations, we developed match-seq, an imaging-free, sequencing-based cell proximity tracing system that uses virus-like particles to transmit barcodes between neighboring cells. Barcoded mRNAs are transmitted from “sender” cells to nearby “receiver” cells, thereby establishing a spatial linkage that can be computationally reconstructed by barcode sequencing after tissue dissociation. We apply this system to an *in vivo* syngeneic murine tumor model, where we observe robust labeling of all immune cell lineages, and leverage barcode labeling to reconstruct cell type niches that recapitulate known tumor spatial biology. Furthermore, we couple this system with CRISPR perturbations and single-cell RNA sequencing to perform genetic screens *in vivo* on tumor-immune interactions at single-cell resolution. We use this method to infer cell-cell spatial relationships and uncover genetic dependencies in cancer cells that change the composition and cell state of their local microenvironments. By calculating a local immune activation signature, we are able to prioritize targets whose deletion enhances anti-tumor immunity through distinct effector cell types: loss of *Tgfb1* engages CD8 T cells and macrophages, *Traf7* loss elicits a CD4 T cell response, and *Nectin3* loss promotes NK cell-mediated immunity, relationships that we validate by *in vivo* immune cell depletion. Notably, *Tgfb1* and *Nectin3* deletion had little effect on cancer cell fitness in the pooled screen yet suppressed tumor growth when deleted throughout the tumor, demonstrating the potential of match-seq to reveal functionally important tumor-immune interactions that would otherwise remain hidden. Together, these results establish a general framework for pooled genetic dissection of cell-cell interactions *in vivo*, extending CRISPR screening from cell-intrinsic phenotypes to the mechanisms by which cells shape and respond to their local microenvironments.

## Introduction

Cell-cell interactions play diverse, complex, and essential roles in shaping tumor progression and outcome. Cancer cells often modulate the tumor microenvironment to favor their own growth and have evolved elaborate strategies to evade immune eradication. Consequently, there is great interest in therapeutic correction of this dysregulated communication, and strategies such as immune checkpoint blockade have proven successful across several cancer types^1–3^. Greater understanding of the complex cell-cell interactions within the tumor microenvironment would not only lead to clarity on resistance mechanisms but also inform the development of new treatments.

While there is a growing appreciation for performing direct measurement of heterogeneous cell-cell interactions^4,5^, tools for doing so remain limited. Pooled CRISPR screens, either in co-culture with immune cells or *in vivo*^6–12^, have been used to infer tumor-immune interactions that alter cancer cell fitness in high-throughput but do not directly measure cell-cell interactions. In contrast, many methods have been developed to label interacting cells, mainly through enzymatic reaction or protein transfer^13–17^, but these protein-based systems are difficult to scale up to measure a large number of heterogeneous interactions. Instead of protein labels, the use of nucleic acid barcode labels would greatly increase the scalability of labeling diverse interactions and has been successfully used with rabies virus in the brain to measure cell-cell connectivity^18–20^.

Here, we developed match-seq, a scalable sequencing-based approach that records spatial proximity between sender and receiver cells through virus-like particle-mediated transfer of barcoded RNA. The use of nucleic acid barcodes provides substantially greater labeling diversity than protein-based approaches and enables cell-cell relationships to be reconstructed by sequencing after tissue dissociation. By combining match-seq with pooled CRISPR screening and single-cell RNA sequencing (Perturb-match), we investigate tumor-immune interactions *in vivo* and simultaneously measure the fitness and transcriptional state of perturbed cancer cells and the composition and state of their local immune microenvironments. This approach enables high-throughput genetic dissection of how cancer cells shape neighboring cells and reveals tumor-immune dependencies that are inaccessible to conventional pooled fitness screens.

## Results

### Recording of spatial relationships using barcoded RNA transfer

To systematically interrogate how cells interact within complex tissues, we sought to develop a scalable approach that could record spatial proximity between genetically defined sender cells and neighboring recipient cells. To do this, we developed match-seq (<u>m</u>ultiplexed <u>a</u>ssociative tagging of <u>c</u>ell interaction <u>h</u>istories by <u>seq</u>uencing), which transfers barcoded mRNA from sender to recipient cells using virus-like particles (VLPs) (**Fig. 1a**). Because immune cells are highly sensitive to viral infection, we sought to identify a VLP that would reduce off-target perturbation due to infection. PEG10 is a retrotransposon-derived protein that forms VLPs packaging mRNA with the *Peg10* untranslated region (UTR) sequences and can be pseudotyped with VSVG to create infectious particles^21^. Because PEG10 is an endogenous protein, we hypothesized that it would evade innate viral sensors and avoid antigenicity due to central tolerance. We compared PEG10 to lentiviral vectors in induced pluripotent stem cell (iPSC) derived macrophages^22,23^, a system that is highly sensitive to viral infection^24^. Infection using integrase-deficient lentivirus (IDLV) induced acute cell death (**Extended Data Fig. 1a, b**), and the cargo mCherry vector was poorly expressed (**Extended Data Fig. 1c**). In contrast, cells infected with PEG10 displayed similar cell morphology compared to infection-naive cells, had significantly reduced cell death, and robustly expressed mCherry rapidly upon addition of PEG10 VLPs (**Extended Data Fig. 1a-c**).

**Figure 1.**
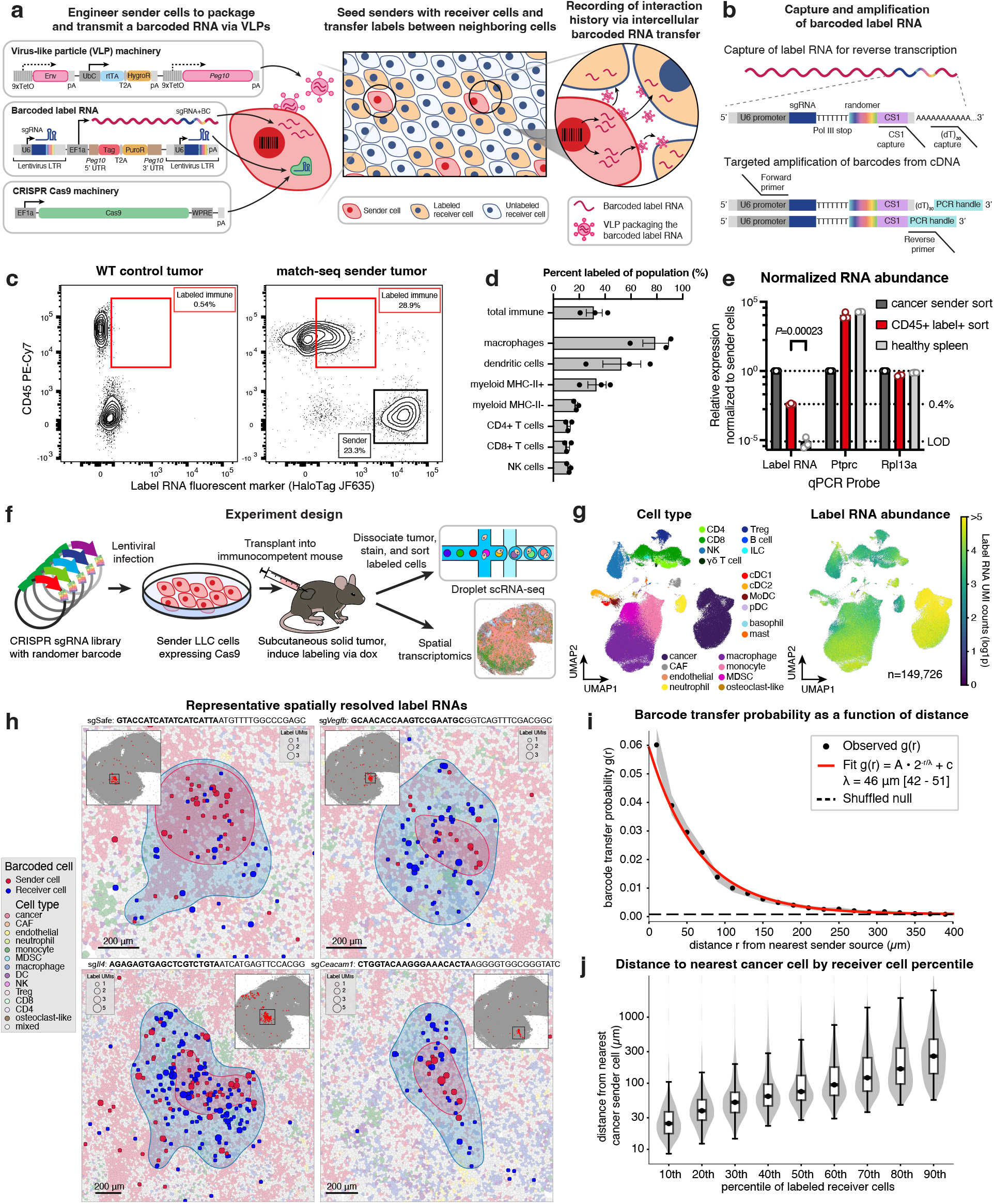
Match-seq labels spatially proximal cells *in vivo* using barcoded RNA transfer. A. Design of match-seq labeling system. Barcoded RNAs are transferred from sender cells to neighboring receiver cells and can be used to reconstruct spatial relationships. B. Reverse transcription of label RNA through poly(dT) and/or capture sequence oligos, followed by targeted PCR amplification, compatible with single-cell RNA-seq. C. Representative flow cytometry of dissociated tumors generated from WT or match-seq LLC sender cells. Cells were stained with anti-CD45 antibody for host immune cells and JF635 HaloTag ligand to detect labeling. D. Percentage of label-positive population for total tumor infiltrating immune cells and different immune cell types, detected by flow cytometry. Data are means of n=3 tumor replicates ± s.e.m. E. Detection of label RNA and control transcripts using RT-qPCR from sorted cancer sender cells, label-positive immune cells, and control cells from a healthy spleen. Values are normalized to the mean of the cancer sender samples. Data are means of n=3 tumor replicates ± s.e.m. *P* value from Welch’s *t*-test. LOD: limit of detection. F. Diagram of Perturb-match experiment. Sender LLC-Cas9 cells were transduced with a library of perturbations and transplanted into syngeneic wildtype mice. Tumors were dissociated, and both cancer cells and label-positive immune cells were sorted for scRNA-seq. Paired samples were also processed using Visium HD 3’ for spatial detection of label RNAs. G. UMAP of scRNA-seq gene expression data, colored by cell type annotations (left) or label RNA UMIs detected (right). H. Representative images of barcodes detected in sender cancer cells (red) and receiver non-cancer cells (blue). sgRNA protospacer sequence is in bold. Shaded contours show kernel density estimates of sender and receiver regions, respectively. Dot size is scaled by the number of label RNA UMIs detected in each cell. All cells positive for the indicated barcode in the tissue are shown in red in the inset panel. I. Probability of barcode transfer as a function of distance. The red line shows an exponential function fit to the observed labeling probability at a given distance. The shaded area shows 95% confidence interval and the dashed line shows expected value of the permuted null distribution. J. Distance of labeled receiver cell from nearest sender cell by percentile of labeled receiver cells.

To perform an unbiased analysis of the transcriptomic effects of each virus on infection, we sorted infection-naive, mCherry^+^ infected, or mCherry^−^ bystander iPSC-derived macrophages in cultures that were either exposed to lentivirus (LV), IDLV, or PEG10 and performed bulk RNA-seq. Principal component analysis (PCA) revealed that cells in the LV and IDLV conditions drove the primary axis of variation compared to PEG10 and infection-naive cells (**Extended Data Fig. 1d**). Both infected and bystander cells in the lentivirus and IDLV conditions had substantial changes in their transcriptomes compared to infection-naive cells, whereas infection with PEG10 displayed markedly fewer significantly differentially expressed genes (10,110 genes, 8,943 genes, and 1,184 genes at adj. *P* < 0.05 in LV, IDLV, and PEG10, respectively) as well as reduced magnitude of gene expression changes compared to infection-naive cells (mean absolute fold change of 3.09, 3.14, and 1.60 of adj. *P* < 0.05 genes for LV, IDLV, and PEG10, respectively) (**Extended Data Fig. 1e, f**; **Supplementary Table 1**). As expected, cells infected with LV or IDLV upregulated genes in pathways related to virus sensing and interferon, while upregulation of these pathways in PEG10-infected cells was reduced (**Extended Data Fig. 1g**). Based on these qualities, we selected PEG10 as the VLP vehicle to deliver barcoded RNA for cell-cell labeling.

To engineer sender cells, we stably integrated a construct encoding PEG10 and vesicular stomatitis virus envelope glycoprotein (VSVG) under tetracycline-responsive promoters as well as a constitutively expressed reverse tetracycline transactivator (rtTA) via DNA recombination using the large serine recombinase Cp36^25^ (**Fig. 1a**). The label RNA cassette was stably integrated at low multiplicity of infection using a lentiviral vector, which consists of a fluorescent marker, the *Peg10* UTR sequences, as well as an sgRNA and a randomer barcode embedded into the lentiviral 3’ long terminal repeat (LTR) (**Fig. 1b**)^26^. The randomer serves as a unique molecular identifier (UMI) to distinguish different sender clones that have the same sgRNA and also increase library diversity^27,28^. The *Peg10* UTR sequences target the RNA transcript for packaging into PEG10 VLPs that are pseudotyped by VSVG and released from the cell as infectious particles^21^. Upon infection, recipient cells express the fluorescent marker encoded on the label RNA and can be enriched by fluorescence-activated cell sorting (FACS). Sequencing of the barcodes in the 3’ LTR can be performed to match spatially proximal sender and receiver cells in a fully sequencing-based manner. Because the label RNA is non-integrating and degrades, our system is capable of detecting recent or transient interactions. Importantly, match-seq is compatible with 3’ scRNA-seq and contains a capture sequence for enhanced detection^29^, enabling measurements at single-cell resolution (**Fig. 1b**). Finally, Cas9 can be expressed in the sender cells to utilize the sgRNA for genetic perturbation screens in the sender cells that alter phenotypes in proximal receiver cells.

To test label transfer *in vivo*, we generated sender cell lines by stably integrating match-seq cassettes into Lewis lung carcinoma (LLC) cells and transducing a label RNA cassette encoding GFP. While match-seq designs that constitutively expressed PEG10 and VSVG resulted in tumor rejection, a tetracycline-controlled system allowed for tumor establishment (**Extended Data Fig. 2a**). Robust induction of PEG10 was improved by addition of the *HBB* IVS2 intron^30^ (**Extended Data Fig. 2b**). GFP sender cells were injected subcutaneously into wildtype syngeneic mice and allowed to establish for 16 days before inducing labeling by doxycycline for 48h. The tumors were then dissected, dissociated, and profiled using antibody staining and flow cytometry to measure GFP labeling activity in the immune compartment. We observed labeling of 3.4% of CD45^+^ cells (mean of n=4 tumors), with GFP detection in both myeloid and lymphoid cell lineages (**Extended Data Fig. 2c, d**). We reasoned that receiver detection could be improved with the use of a brighter fluorescent probe. To this end, we replaced the fluorescent protein in the label RNA with HaloTag^31^ and detected labeling using Janelia Fluor 635 HaloTag ligand (JF635-HTL), a dye that fluoresces brightly upon reaction with HaloTag^32^. Detection of HaloTag receivers using JF635-HTL improved labeling detection to 32% of CD45^+^ receiver cells (mean of n=3 tumors), with efficient labeling of both myeloid and lymphoid lineage cell types (**Fig. 1c, d**). Transfer of the label RNA to recipient immune cells was detected by sorting CD45^+^ JF635^+^ cells and performing RT-qPCR. The label RNA was detected at 0.4% of the abundance in sender cells and was reliably detected above negative control healthy spleen samples (*P*=0.00023; Welch’s *t*-test) (**Fig. 1e**).

To demonstrate the scalability of barcoding, we generated match-seq LLC cells barcoded with a library of 1000 safe control sgRNAs with a 16 base randomer in order to create a high diversity library. Cells were injected subcutaneously into wildtype mice, and sender LLC cells and receiver CD45^+^ cells were sorted after tumor dissociation. RNA fractions were then collected from both sender and receiver cell pellets, and genomic DNA (gDNA) was additionally collected from sender cells in order to estimate the abundance of senders of each barcode. We detected 7,115 unique barcodes across all three replicates, and gDNA abundance was well correlated with barcode abundance in the RNA fraction of both sender cells and receiver cells (R=0.76 and 0.65, respectively), suggesting that labeling rate is proportional to sender abundance (**Extended Data Fig. 2e**). To demonstrate that match-seq can be applied to other tumor models and contexts and visualize spatial spread of label RNAs, we engineered murine 4T1 breast cancer cells with match-seq and established lung metastases through tail vein injection. Label RNAs were detected using RNA fluorescence *in situ* hybridization (FISH), and we observed label RNAs in host immune cells that were adjacent to sender cancer cells (**Extended Data Fig. 2f**).

### Match-seq labels spatially proximal cells at single-cell resolution

To demonstrate the utility of match-seq, we applied it in combination with a CRISPR KO perturbation library (Perturb-match) in the sender cells to investigate the effect of genetic perturbations on cancer cells and their local microenvironments *in vivo* (**Fig. 1f**). We designed a bespoke library consisting of 417 sgRNAs targeting 132 expressed genes in LLC cells that are involved in extracellular signaling and sensing of key immune signaling pathways, such as interferon, TGFB superfamily, and TNF superfamily members (**Supplementary Table 2**; **Methods**). Cells were subcutaneously engrafted into wildtype immunocompetent mice and allowed to establish for 11 days before induction of labeling for 48h. Nine tumors were dissociated and sample-hashed using MULTI-seq^33^ before pooling and FACS for label-positive immune cell populations and cancer sender cells for scRNA-seq library construction (**Methods**). FACS was used to enrich for T, NK, and dendritic cells, as these populations were present in lower frequencies (**Extended Data Fig. 3a**). In parallel, tissue from one of the tumors was processed using Visium HD 3’ spatial transcriptomics (ST) in order to assess ground truth spatial measurements.

After quality control and multiplet removal, the Perturb-match dataset consisted of 149,726 single cell transcriptomes, which were annotated with cell types and cell states based on marker gene expression (**Fig. 1g**; **Extended Data Fig. 4a-h**; **Supplementary Table 3**). Cells were robustly assigned to tumor replicates using MULTI-seq barcodes (**Extended Data Fig. 4i**). Label RNAs were captured using both poly(dT) and CS1 capture (**Extended Data Fig. 5a**). We detected label RNAs across all major cell lineages, with comparable labeling across immune cell types and higher labeling in fibroblast and endothelial cells (**Extended Data Fig. 5b**). In general, cells received more than one barcode, with a median of 7 unique barcodes per cell across cell types (**Extended Data Fig. 5c**). We recovered 410,703 unique barcodes across nine tumors and a mean of 5.83 receiver cells labeled per unique barcode (**Extended Data Fig. 5d, e**). sgRNA abundances were well correlated across tumor replicates, suggesting good reproducibility (**Extended Data Fig. 5f**). As we observed with the bulk measurements, the proportion of labeled receiver cells was well correlated with the proportion of corresponding sender cells for that barcode (R=0.75) with a proportional increase between percentage of senders and labeled receivers (slope=0.96) (**Extended Data Fig. 5g**).

To confirm that label transfer was spatially proximal, we assessed the spatial distribution of label barcodes in a tumor sample processed using spatial transcriptomics. A tissue section from one of the tumors that was processed for scRNA-seq was additionally profiled using Visium HD 3’, which allowed for spatially resolved measurements of gene expression and label RNA barcodes at a subcellular 2 µm bin resolution. After cell segmentation, cell types were assigned using the scRNA-seq dataset as a reference^34^, and we observed separation of cell types in expression of known marker genes (**Extended Data Fig. 6a-c**). We then mapped label RNA barcodes to either sender cancer cells or receiver cells at single-cell resolution, using the randomer to distinguish sender clones with the same sgRNA perturbation. Label RNAs were well detected across the entire tissue sample (**Extended Data Fig. 6d**). By examining the spatial distribution of individual barcodes, we found strong evidence that barcodes were spatially localized rather than randomly distributed (*P* < 0.05 for 578/601 barcodes (96.2%), permutation test) (**Extended Data Fig. 6e**; **Supplementary Table 4**), which has been observed in other studies with spatially resolved barcoding of solid tumors^35–37^. In many cases, we were able to identify a well-defined spatial region with detection of a unique barcode in several cells, often containing a subregion of cancer sender cells (**Fig. 1h**; **Supplementary Table 4**). Matching receiver cells with the nearest sender cell, we observed that the probability of barcode transfer decayed exponentially with distance, with the labeling probability halving every 46 µm (**Fig. 1i**). Across all barcodes, the median distance of the 90th percentile of labeled receiver cells was within 250 µm from the nearest sender cell (**Fig. 1j**). These data support that match-seq can be used to label spatially proximal cells *in vivo*.

Transcript capture efficiency in the Visium ST dataset was relatively low; in our datasets, there were approximately 20-fold fewer UMIs and 7-fold fewer genes detected in the ST data than in the scRNA-seq (**Extended Data Fig. 6f**). While the ST dataset was limited to a 2-dimensional tissue section, match-seq was able to enrich and stochastically sample cell-cell interactions from the entire tissue. Thus, match-seq was able to sample many more spatial neighborhoods (median sgRNA coverage of 38 vs 1) (**Extended Data Fig. 6g**) and could also be coupled with FACS to enrich for rarer cell types, such as T cells and NK cells (**Extended Data Fig. 3a**).

### Genetic perturbations induce changes in cancer cell fitness and local microenvironment

To use match-seq to query the spatial ecology of the tumor microenvironment, we examined the cell types associated with each barcode, with the model that cells carrying the same barcode reflect spatially proximal cells. Grouping barcodes with similar cell type compositions, we recovered families of barcodes that were enriched for distinct combinations of cell types. For example, we observed a TAM enriched cluster containing barcodes shared among macrophages and Tregs, as well as several immune-rich clusters enriched for diverse lymphocytes and dendritic cells (**Fig. 2a**). Deeper analysis of consistently co-occurring cell types across tumor replicates revealed four major groups: (1) a stem-like immune hub containing stem-like CD4 and CD8 T cells, (2) an immune rich cluster with effector lymphoid populations and monocytes, (3) a tumor-associated immune group with enrichment for macrophages and antigen-presenting cells, and (4) a tumor-core with mostly cancer cells (**Fig. 2b**). Leveraging the high resolution of scRNA-seq data, we were able to discriminate cell states within cell types that had differential co-localization patterns (**Extended Data Fig. 4b-h**; **Supplementary Table 3**); for example, proliferative macrophages preferentially associated with tumor cells, while *Nos2*^high^, *Ccr2*^high^, and activated macrophages were more likely to be found within immune-rich niches. Several of these relationships closely recapitulated spatial programs observed in human tumors, including the association of SPP1^high^ macrophages with hypoxic tumor regions^38^ and the enrichment of CCL8^high^ macrophages within the tumor core^39^. These broad groupings are consistent with the paradigm of immune-rich, immune-excluded, and immune-desert spatial archetypes^40–42^ as well as previous reports of co-localization of stem-like immune hubs^43^. These results illustrate the ability of match-seq to recover spatial relationships using barcode matching that recapitulate biologically conserved features of tumor organization without direct spatial imaging.

**Figure 2.**
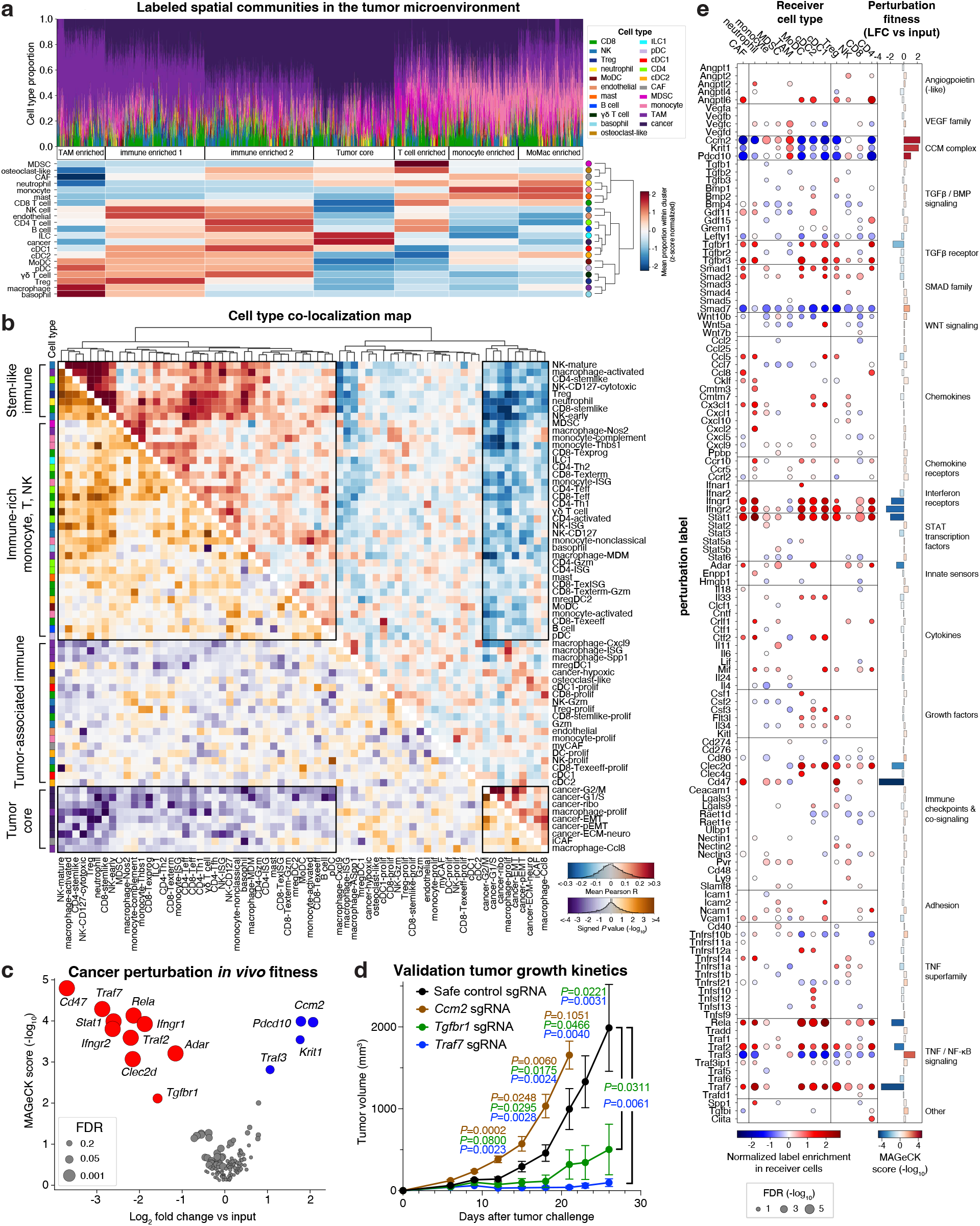
Perturb-match maps cancer perturbation spatial communities. A. (Top) stacked barplot showing cell types labeled by a unique label RNA barcode, ordered by *k*-means clustering. Downsampled using geosketch to n=5000 barcodes. (Bottom) Row-normalized mean proportion for each cell type within *k*-means clusters. B. Co-localization map showing spatially correlated cell types across tumor replicates. Upper triangle shows mean Pearson correlation coefficient of cell types across spatial communities across tumor replicates, as defined by community detection using the barcode graph. Lower triangle shows two-tailed *P* values (signed -log_10_) calculated using Student’s *t*-test. Frequency histograms are inset in color bars. Colored boxes on the left indicate cell type. Heatmap is ordered on hierarchical clustering on Pearson correlation and boxes indicate immune-rich (top left) and tumor-core (bottom right) clusters, and their anti-correlation (top right and bottom left). Cell type colors correspond to legend in (A). n=9 tumor replicates. C. Volcano plot showing perturbation effects on cancer cell fitness compared to input sgRNA abundance. Calculated from cancer cell counts from the Perturb-match dataset. Red points are significantly depleted; blue points are significantly enriched (FDR < 0.1). Effect size is the mean log_2_ fold change and significance is calculated using MAGeCK RRA. n=9 tumor replicates. D. Tumor growth kinetics of *Ccm2*-null, *Tgfbr1*-null, *Traf7*-null, or safe control sgRNA LLC tumor in wildtype syngeneic mice. n=5 (*Ccm2*, *Tgfbr1*, *Traf7*) or n=10 (safe) mice per group. Data are mean ± s.e.m. *P* values from Student’s *t*-test. E. Dotplot shows enrichment or depletion of labeling by different tumor perturbations in neighboring receiver cell types, relative to cancer sender cell abundance and normalized to safe control sgRNA labeling distribution. Perturbations are grouped by gene target family. Dot color represents effect size and dot size represents FDR. Only points with FDR < 0.1 are shown. Statistical significance calculated using RRA. Bar plot shows perturbation fitness, as calculated in C.

To identify genetic perturbations that increased or decreased cancer cell fitness *in vivo,* cancer cells in the dataset were assigned to an sgRNA barcode, and an analysis of the enrichment of sgRNAs compared to the input population was performed (**Fig. 2c**; **Supplementary Tables 5, 6**). For perturbations that caused significant changes in cancer cell fitness *in vivo*, we observed agreement between different sgRNAs targeting the same gene, as expected (**Extended Data Fig. 7a**). Changes in *in vivo* sgRNA representation were largely due to selection pressures from the tumor environment, rather than changes in proliferation that could be observed *in vitro* (**Extended Data Fig. 7b**). The most enriched targets were members of the cerebral cavernous malformation (CCM) complex, *Krit1*, *Ccm2*, and *Pdcd10* (false discovery rate (FDR) < 0.03), which were previously found to be enriched in an *in vivo* screen of the LLC cell line dependent on immune selective pressure^6^. Perturbations that reduced cancer cell fitness included previously reported immune checkpoints *Cd47*^44^ and *Adar*^45^, as well as genes involved in IFNγ signaling (*Ifngr1*, *Ifngr2*, *Stat1*)^6,46,47^, and NF-κB signaling (*Rela*, *Traf2*, *Traf7*)^48^ (FDR < 0.00001) (**Fig. 2c**; **Extended Data Fig. 7a**; **Supplementary Table 6**).

We individually validated *Ccm2* as a perturbation that conferred a fitness advantage and *Tgfbr1* and *Traf7* as perturbations that conferred reduced tumor growth *in vivo* compared to safe control sgRNA tumors (**Fig. 2d**). Although TGFβ receptors can function as tumor suppressors^37^, in our tumor model, loss of *Tgfbr1* reduced tumor growth, suggesting a context-dependent role for TGFβ sensing in immune evasion. TRAF7 is a non-canonical member of the TNF receptor-associated factor (TRAF) family: while it contains N-terminal RING finger and zinc finger domains, similar to other members in the family, it has WD40 domain repeats instead of the conserved C-terminal protein binding domain found in other TRAF proteins^49^. TRAF7 has been reported to act in both pro- and anti-tumor contexts^50–52^, and although it is not mechanistically understood how loss of *Traf7* conferred sensitivity to immune pressure in our screen, we were able to resolve how *Traf7*-null cells remodeled their local microenvironment using match-seq.

To investigate how genetic perturbations in cancer cells changed interaction frequencies with different immune cell types, we examined the enrichment of sgRNA labels within different receiver cell types, normalized to the cancer cell sgRNA abundance and the safe control sgRNA labeling distribution (**Fig. 2e**; **Supplementary Tables 7, 8**; **Methods**). Broadly, perturbations that increased cancer cell fitness, such as knock out of CCM complex members, reduced interactions with lymphocytes while increasing interactions with macrophages, MDSCs, and monocytes (FDR < 0.001), as might be expected for perturbations that confer resistance to immune pressure. Conversely, perturbations that reduced cell fitness tended to increase interactions with lymphocytes and dendritic cells, such as *Traf7*, *Rela*, *Clec2d*, *Ifngr1*, and *Ifngr2* (FDR < 0.001), reflecting a “hot” immune environment.

### Genetic perturbations induce changes in cancer cell transcriptional states

To assess how the genetic perturbations changed the transcriptomes of cancer cells, we performed differential gene expression analysis (**Supplementary Table 9**) followed by consensus non-negative matrix factorization (cNMF) to group perturbations that had similar gene expression changes^53^. cNMF yielded 15 factors, and we recovered factors that represented genes in related signaling pathways, such as type I and type II interferon, CCM complex, NF-κB signaling, and vascular endothelial growth factor members (**Fig. 3a**; **Supplementary Table 10**). Pathway enrichment analysis of the factor gene loadings showed concomitant pathway regulation changes (**Supplementary Table 11**). For instance, knock out of NMF factor 1 genes *Ifngr1*, *Ifngr2*, and *Stat1* resulted in downregulation of response to type II interferon, while knock out of NMF factor 2 genes *Ifnar2* and *Stat2* resulted in downregulation of response to type I interferon. CCM complex genes in factor 15 were associated with upregulation of actin remodeling, consistent with a recent study on the CCM complex in endothelial cells^54^.

**Figure 3.**
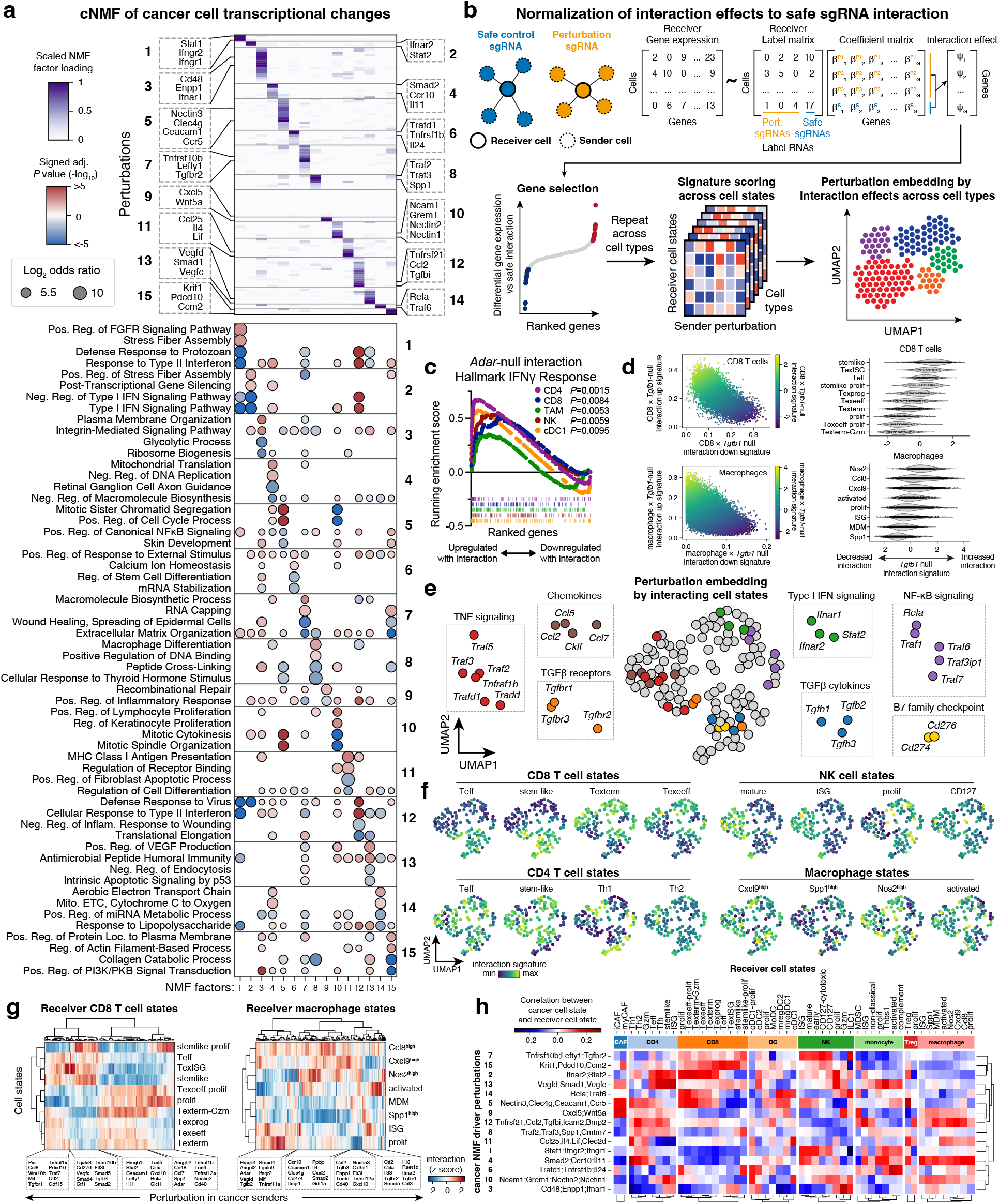
Perturb-match measures how cancer cell genetic perturbations change cell intrinsic programs and paired local microenvironment responses. A. NMF factorization of cancer transcriptome changes associated with genetic perturbations. Top heatmap shows scaled factor loadings for perturbations and bottom dot plot for Gene Ontology pathway enrichment on gene loadings. Dot size represents log_2_ odds ratio and dot color represents signed -log_10_ adjusted *P* value. B. Diagram of calculation of interaction effects on receiver cells. Top shows modeling of safe sgRNA and perturbation sgRNA labeling on receiver cell gene expression. Bottom shows signature creation from differential gene expression, mapping to cell states, and perturbation embedding. C. GSEA for Hallmark Interferon Gamma Response gene set for differentially expressed genes in various receiver cell types comparing interaction with *Adar*-null vs safe control cancer sender cells. D. Single cell scoring of interaction signatures with *Tgfb1*-null cancer cells in CD8 T cell and macrophages, colored by summary interaction score. Scores are plotted by cell state on the right. E. UMAP embedding of perturbations based on cell state interaction scores. Groups of perturbations in TNF signaling, chemokines, TGFβ receptors, TGFβ cytokines, IFNα signaling, NF-κB signaling, and B7 family checkpoint ligands are highlighted. F. Perturbation UMAPs colored by median cell state interaction scores, grouped by cell type. ISG: interferon-stimulated gene; Teff: effector-like T cell; Texterm: terminally exhausted T cell; Texeeff: early effector exhausted T cell. G. Heatmaps showing median interaction scores for cell states in CD8 T cells and macrophages. H. Correlation of cancer NMF cell states (as in A) and receiver cell states (as scored using method described in B).

### Cancer cell perturbations influence local microenvironmental cell state

Because receiver cells often contained a mixture of barcodes belonging to different perturbations (**Extended Data Fig. 5c**), we developed a computational framework to control for baseline interaction effects and estimate how genetic perturbations in cancer cells affect transcriptional state in receiver cells relative to interaction with a safe control sgRNA cancer cell (**Fig. 3b**; **Methods**). Since we observed labeling rate to decay exponentially with distance from a sender cell (**Fig. 1i**), we treated label RNA UMIs as a quantitative measure of proximity and fitted regression models to estimate how interaction with a certain perturbation differentially regulates gene expression relative to safe control sgRNAs (**Supplementary Table 12**). Using this method we were able to recapitulate previously reported results, such as upregulation of IFNγ response in cells proximal to *Adar*-null cancer cells across several cell types^45^ (*P* < 0.01) (**Fig. 3c**).

For each cell type and cancer cell perturbation, we then derived a gene expression signature from interacting receiver cells and used it to score how closely individual cells resembled that interaction-associated cell state (**Supplementary Tables 13, 14**). As expected, signatures for increased and decreased interaction were negatively correlated and could be projected onto single cells to look for signature enrichment in different cell states (**Fig. 3d**). For example, the signature for CD8 T cell interaction with *Tgfb1*-null cancer cells showed enrichment in stem-like, interferon-stimulated gene exhausted T cells (TexISG), and effector-like (Teff) cell states with reduced scores in exhausted T cell populations. In macrophages, *Tgfb1*-null interaction signature was enriched in pro-inflammatory macrophage states (*Nos2*^high^, *Ccl8*^high^, *Cxcl9*^high^) and reduced in suppressive states (*Spp1*^high^, *Ccr2*^high^), again suggesting the presence of a productive immune response neighboring *Tgfb1*-null cancer cells. UMAP embedding of perturbations on the basis of their interaction profiles across all interacting cell states showed clustering of genes in similar pathways, such as chemokines, TGFβ signaling, TNF sensing, and NF-κB signaling (**Fig. 3e**; **Extended Data Fig. 8a**). Overlaying the cell state score on the embedding revealed how different perturbations enrich for different cell state neighborhoods (**Fig. 3f**; **Extended Data Fig. 8b-f**). Loss of genes encoding TGFβ isoforms and B7 family immune checkpoints resulted in increased interaction with stem-like and effector CD4 and CD8 populations, consistent with the known role of these factors as immunosuppressive^55–60^. In contrast, loss of genes encoding chemokines involved in myeloid cell recruitment (*Ccl2*, *Ccl5*, *Ccl7*) resulted in association with *Cxcl9*^high^ and *Nos2*^high^ macrophages, which have been described as pro-inflammatory macrophage states^61^. Tumor cell secretion of these chemokines has been described to recruit immunosuppressive myeloid populations^62–65^. We found enrichment of an IFNγ response signature in *Ccl2*-null cancer cells (**Fig. 3a**, **Supplementary Tables 9-11**), suggesting that KO of these genes results in a pro-inflammatory local environment.

Interestingly, we observed clustering of perturbations involved in TNF sensing (*Tnfrsf1b*, *Tradd*, *Traf2*, *Traf3*, *Traf5*), which were observed to enrich for interactions with mature and ISG NK cell populations, while perturbations associated with the canonical NF-κB pathway (*Rela*, *Traf1*, *Traf6*, *Traf7*) clustered separately with a different niche enriched for Th1 CD4 T cells. Similar distinction between TRAF family members was also observed by gene expression changes in the cancer cell transcriptomes (**Fig. 3a**; **Supplementary Table 10**), showing that incorporating neighborhood remodeling offers complementary views to perturbation effects. Clustering perturbations based on their cell state interaction profiles revealed groups of perturbations that have similar effects on cell types (**Fig. 3g**; **Extended Data Fig 8g-i**). For instance, we observed four broad groups of perturbations that influenced CD8 T cell states to stem-like / effector, early exhaustion, terminal exhaustion, or proliferative phenotypes.

Having estimated the effects of perturbations in both cancer cells as well as their neighboring immune cells, we reasoned that we could leverage the diversity of transcriptional programs created by the perturbation library to understand how cancer cell states were related to their local microenvironments. To this end, we correlated the observed transcriptional heterogeneity between sender and receiver cells across perturbations (**Fig. 3h**). Strikingly, this analysis recovered expected biological results, such as reduced CD8 T cell interaction but increased NK cell interaction with NMF factor 1 (cancer cells with knockout of *Stat1*, *Ifngr1*, *Ifngr2*), consistent with the notion that IFNγ upregulates MHC-I, which is required for T cell cytotoxicity but regulates NK cells through missing-self. Indeed, MHC-I genes are among the most downregulated genes in NMF factor 1 (**Supplementary Tables 9, 10**).

### Modeling of local immune activation unmasks distinct cancer immune dependencies from pooled screening

Identification of cancer targets that would activate neighboring immune cells could have potential therapeutic value but could be difficult to detect in conventional pooled screens that solely measure growth. To find such factors, we developed a score to identify perturbations that preferentially increase local immune activation (LIA) (**Fig. 4a**). In brief, differential gene expression changes upon interaction for each cell type were factorized by cNMF to derive factors expressing groups of perturbations that had similar effects on each cell type (**Supplementary Table 15**). Then, a factor corresponding to immune activation was identified for each cell type (**Extended Data Fig. 9a**), and the mean perturbation loadings for these factors across cell types was used to derive a LIA score for each perturbation (**Supplementary Table 16**; **Methods**). Plotting the LIA score against the observed cancer cell fitness highlighted a subset of perturbations that had no effect on fitness in the pool but were nonetheless predicted to activate neighboring immune cells (**Fig. 4b**).

**Figure 4.**
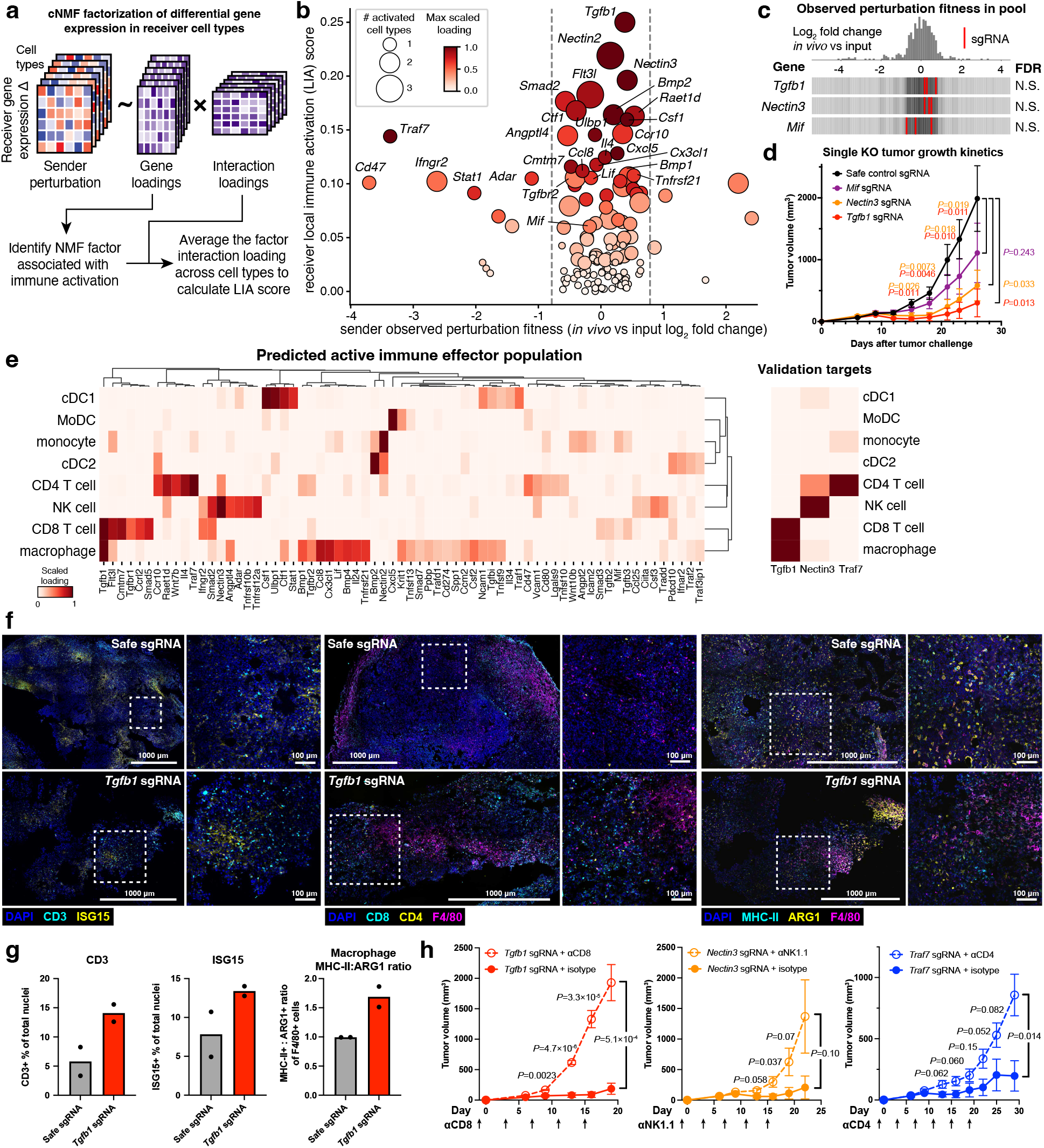
A local immune activation signature nominates novel cancer dependencies that are masked by conventional pooled screening. A. Diagram of method to calculate local immune activation (LIA) score of perturbations. B. Scatter plot of perturbations comparing impact on cancer cell fitness versus LIA score. Dot size represents the number of immune cell types with >0.2 scaled factor loading, and dot color is the max scaled factor loading across all effector cell types. Dashed lines show 1 standard deviation. C. Distributions of sgRNA abundance of cancer cells with *Tgfb1*, *Nectin3*, and *Mif*, normalized to the input sgRNA abundance. N.S.: not significant (FDR > 0.1). D. Tumor growth kinetics of *Mif*-null, *Nectin3*-null, *Tgfb1*-null, or safe control sgRNA LLC tumors in wildtype syngeneic mice. n=4 (*Tgfb1*), n=5 (*Mif*, *Nectin3*), or n=10 (safe) mice per group. Data are mean ± s.e.m. *P* values from Student’s *t*-test. E. Prediction of active immune effector population for cancer perturbations based on local immune activation score. Validation targets *Tgfb1*, *Nectin3*, and *Traf7* are re-plotted on the right. F. Representative images of safe control sgRNA and *Tgfb1*-null tumors stained with DAPI, CD3 and ISG15 (left), DAPI, CD8, CD4, and F4/80 (middle), and DAPI, MHC-II, ARG1, and F4/80 (right). Boxed regions are displayed to the right of their respective images. G. Quantification of microscopy of CD3^+^ of total cells (left), and ISG15^+^ of total cells (middle), and MHC-II^+^:ARG1^+^ ratio of F4/80^+^ cells (right). Data are means of n=2 tumor replicates. H. Tumor growth kinetics for LLC cells with the indicated gene knockout and cell depletion treatment. Arrows indicate treatment timing. Data are mean ± s.e.m. n=5 mice per group. *P* values from Student’s *t*-test.

Among these perturbations were several genes that have previously been described to have immunosuppressive functions. The gene with the highest LIA score, *Tgfb1*, has been reported to facilitate immune exclusion in solid tumors^60^ and is known to suppress cytotoxic T cell^66^ and macrophage activation^67,68^. Other perturbations with high LIA score include the genes encoding TIGIT ligands NECTIN2 (CD112) and NECTIN3 (CD113)^69,70^, macrophage growth factor M-CSF^71,72^, and cytokines IL-4^2,73^ and LIF^74,75^, which have been described to have immunosuppressive properties. Although FLT3L is not typically studied in the context of cancer cell expression, we observed that loss of *Flt3l* in cancer cells resulted in a high LIA score. This is corroborated by a previous study reporting loss of *Flt3l* in cancer cells resulted in reduced Treg licensing and an enhanced adaptive immune response^76^. Interestingly, several genes encoding secreted factors had high LIA scores, including *Angplt4*, *Cx3cl1*, *Ctf1*, and *Cmtm7*. While the described roles of these factors in tumor immunity literature have been inconsistent^77–81^, our analysis nominates them as putative immunosuppressive factors in the LLC tumor context.

We hypothesized that certain perturbations with a high LIA score, such as *Tgfb1*, may not exhibit an observable fitness effect in a pooled CRISPR screen setting, where neighboring cells in the mosaic environment may be able to provide sufficient quantities of the factor to mitigate any overall growth phenotypes. However, we reasoned that single-gene knockout tumors for these genes would lead to immune activation throughout the tumor and lead to immune-mediated tumor suppression. To test this, we generated knockout cell lines for *Tgfb1*, *Nectin3*, and *Mif*, which have high to moderate LIA scores. We confirmed that sgRNAs targeting these genes had no cancer fitness effect in our pooled screen dataset (**Fig. 4c**), or in published *in vivo* pooled CRISPR screens in LLC cells, which included an immunodeficient NSG *in vivo* control group, where they were also found to have no fitness effect^6^ (**Extended Data Fig. 9b**). Thus, these genes could not have been discovered in a conventional pooled CRISPR growth screen. Compared to cells carrying safe control sgRNA, *Tgfb1* and *Nectin3* knockout significantly impeded tumor growth (*P* < 0.05, Student’s *t*-test), while *Mif* knockout modestly reduced tumor burden (**Fig. 4d**).

Using the Perturb-match data, we were also able to predict which immune effector cell population(s) were responsible for driving high LIA scores for each cancer cell perturbation (**Fig. 4e**; **Supplementary Table 16**). While CD8 T cells and macrophages were predicted to drive response to *Tgfb1*-null tumors, CD4 T cells were predicted to act on *Traf7*-null cells, and NK cells were predicted to act on *Nectin3*-null cells (**Fig. 4e**). Immunofluorescence microscopy of *Tgfb1*-null tumors revealed evidence of elevated T cell and macrophage activity, such as increased T cell infiltration, an increased ratio of MHC-II:ARG1-positive macrophages, and increased ISG15 positive cells, compared to safe control sgRNA tumors (**Fig. 4f, g**). Finally, we performed *in vivo* cell depletion experiments for each of the three predicted perturbation-effector cell type pairs, and confirmed that the respective immune cell type was necessary for sustaining an effective immune response to each particular gene knock out (**Fig. 4h**). Although nectin family proteins have been described to inhibit T and NK cells through binding to TIGIT, prior studies primarily point to PVR and CD112 as the highest affinity ligands^11,69,70^. Nonetheless, our data indicates that CD113 restrains NK-mediated immunity, suggesting that factors beyond binding affinity may be at play *in vivo*. In addition, we are able to use match-seq to tie the observed reduced fitness of *Traf7*-null cells *in vivo* to a CD4 T cell response. Although the role of TRAF7 in tumor immunity is relatively obscure, our data suggests that it may play a role in sustaining canonical NF-κB response in this context, given the similar local environment profile of cancer cells carrying *Rela*, *Traf1*, and *Traf6* perturbations (**Fig. 3e**). Overall, this analysis highlights the ability of Perturb-match to recover cancer immune dependencies that can be masked by pooled CRISPR screening for barcode enrichment phenotypes and validates predictions on the active immune population made from the Perturb-match data.

## Discussion

Here, we establish match-seq as a scalable framework for genetically dissecting cell-cell interactions *in vivo*. By coupling VLP-mediated transfer of barcoded RNA with pooled CRISPR screening and single-cell RNA sequencing, Perturb-match simultaneously measures genetic perturbations in sender cells and the identity and transcriptional state of their neighboring receiver cells. We applied this to an immunocompetent tumor model, and could reconstruct known and novel spatially organized cellular communities. Perturb-match revealed how cancer cell perturbations remodel the composition and state of their local immune microenvironments, and uncovered tumor dependencies that were not apparent from cancer cell fitness alone. Importantly, these interaction phenotypes identified not only perturbations that promote local immune activation but also the immune cell populations associated with each response. Together, these findings extend pooled genetic screening beyond cell-intrinsic phenotypes to the high-throughput interrogation of how cells shape and respond to their local environments.

Match-seq offers several advances over existing technologies that trace cell-cell interactions. First, whereas current tools rely on protein transfer or enzymatic labeling^13–17^, genetically encoded RNA barcodes provide substantially greater labeling diversity and can be read out together with the transcriptome by sequencing. We demonstrate that our system is well suited for use in combination with CRISPR pooled screens as well as single-cell RNA sequencing. Second, VLP-mediated transfer eliminates the need for engineering of bespoke synthetic receptors on receiver cells, allowing endogenous host cells to be profiled in wildtype animals. This property was particularly useful in this study, where we performed tumor transplant experiments in wildtype syngeneic mice and observed robust labeling of diverse infiltrating host immune cells. Finally, the discovery that PEG10 VLPs exhibit markedly reduced toxicity and antiviral response in infected cells compared to lentivirus makes this system particularly useful for preserving the immune cell states that the system is intended to study.

Conventional pooled CRISPR screens typically identify genes that change the abundance of the perturbed cell. However, oncogenes and tumor suppressors can have strong effects on the surrounding immune microenvironment without producing a detectable fitness phenotype in a mosaic pooled screen. By identifying perturbations associated with local immune activation states, we recovered several such genes, including *Tgfb1* and *Nectin3*, whose deletion had little effect on representation in the pooled screen but suppressed tumor growth when deleted throughout the tumor. The match-seq interaction data further predicted distinct immune effectors associated with each cancer perturbation, including CD8 T cells and macrophages for *Tgfb1*, CD4 T cells for *Traf7*, and NK cells for *Nectin3*, which we validated by *in vivo* immune cell depletion experiments. These results illustrate how profiling the state of neighboring cells can reveal biologically important cancer dependencies that would otherwise be masked in pooled fitness screens and, more broadly, provide a strategy for prioritizing perturbations according to the type of local immune response they elicit.

Although there are existing methods to measure spatial effects of genetic perturbations using spatial transcriptomics (ST)^36,37,82–84^, we believe that match-seq is highly complementary and offers several important advantages. In comparison with ST, match-seq benefits from greatly increased transcript detection, owing to the maturity of droplet-based scRNA-seq chemistries over nascent ST methods. Furthermore, match-seq is able to sample cells from the entire tissue sample with the option of enriching for rarer cell types through FACS, rather than having to only measure a single 2-dimensional tissue section. Especially in the tumor immunology field, it is increasingly recognized that sampling of diverse spatial regions is required for a complete understanding of the underlying biology^85,86^. Finally, match-seq is able to leverage existing technologies that have been developed for scRNA-seq assays. For instance, we used MULTI-seq to pool cells from multiple tumor replicates to increase statistical power for downstream analyses^33^. Future optimization of VLP tropism, labeling duration, and label RNA stability may tune the cellular and temporal range over which interactions are recorded, potentially enabling recording of dynamic cellular relationships and migration, which is an advantage of cell labeling systems over microscopy based approaches.

While we focused here on tumor-immune interactions, the underlying match-seq framework is not restricted to cancer or to immune activation as an endpoint. Interaction profiles could be used to prioritize perturbations that alter essentially any measurable receiver-cell phenotype, including differentiation, stress, tissue repair, signaling, or other multicellular behaviors. Match-seq could also be combined with larger perturbation libraries and emerging single-cell multi-omic measurements to connect sender cell genotypes with increasingly rich descriptions of neighboring cell states. More broadly, by measuring the local consequences of genetic perturbations at scale, match-seq presents a strategy to systematically define the functional regulators of multicellular interactions in cancer and other biological contexts.

CRISPR screening has demonstrated great utility in identifying genetic regulators of cell intrinsic functions and pathways. Our approach extends pooled CRISPR screens to explicitly measuring intercellular effects. We anticipate that match-seq can be broadly applied to a variety of biological systems to systematically interrogate the spatial consequences of genetic perturbations.

## Methods

### Molecular cloning

Plasmids were cloned using Gibson Assembly with ET SSB^88^ (NEB M2401S) and verified using whole plasmid sequencing performed by Plasmidsaurus using Oxford Nanopore Technology with custom analysis and annotation. PCRs were performed using CloneAmp HiFi Premix (Takara 639298). mmPEG10rc4 was PCR amplified from pCMV-mmPEG10rc4 (Addgene 174858), *Peg10* UTR sequences were PCR amplified from pCMV-mm.cargo(Cre) (Addgene 174862), VSVG was PCR amplified from pCMV-VSV-G (Addgene 8454), and *HBB* IVS2 was PCR amplified from pSA073 (Addgene 249134).

A randomer barcode template oligo (gPD041; **Supplementary Table 17**) was ordered from Integrated DNA Technologies (IDT), and 0.15 pmol of ssDNA template was PCR amplified for 14 cycles using KAPA HiFi HotStart ReadyMix (Roche KK2602) and gel purified on a 2% TBE gel. PCR products were mixed with 500 ng of backbone vectors pPD652 (Addgene XX) at a 10:1 molar ratio, reacted using 60 cycles (37°C to 16°C, 1 min each) of PaqCI Golden Gate cloning in a 15 μL reaction, followed by a 5 min 60°C end soak (NEB R0745; NEB M1100). This reaction was then directly digested with 2 μL of BlpI in a 100 μL reaction with rCutSmart buffer at 37°C for 90 min to cut unreacted backbone plasmids (NEB R0585). Ligation products were column purified using MinElute columns (Qiagen 28006) and eluted with 10 μL of elution buffer (EB; Qiagen). Two μL of ligation products were mixed with 50 μL of Endura electrocompetent cells (Lucigen 60242-2) and electroporated following the manufacturer’s instructions using a Gene Pulser Xcell (BioRad). Cells were immediately recovered in 4 mL of recovery media, shaking at 250 rpm at 30°C for 2h and then directly inoculated into 250 mL of terrific broth containing 100 μg mL^−1^ carbenicillin (RPI 50-213-248). Liquid cultures were shaken at 250 rpm at 30°C for 16h before plasmid extraction (Qiagen 12663).

sgRNA libraries were synthesized (Twist Bioscience), PCR amplified for 14 cycles using Herculase II (Agilent 600679), and gel purified on a 2% TBE gel. The randomer backbone plasmid was pre-digested with BlpI and Esp3I (NEB R0585; R0734) for 3h at 37°C and gel purified on a 1% TAE gel. PCR products were mixed with 300 ng of digested backbone plasmid at a 10:1 molar ratio, digested for 10 min at 42°C, and reacted using 60 cycles (42°C to 16°C, 5 min each) of BsmBI-v2 Golden Gate cloning in a 15 μL reaction, followed by a 30 min 42°C final digestion and 5 min 60°C end soak (NEB E1602). Ligation products were column purified, amplified, and extracted, as described above.

### Plasmid list

pPD430: match-seq machinery vector

pPD652: match-seq label RNA lentiviral construct cloning backbone

pPD665: pPD652 with 16 nt randomer barcode

pPD666: pPD665 with libPD006 KO sgRNA library

pPD670: Cas9-T2A-BFP-P2A-BSD lentiviral construct

pPD164: tetracycline-inducible PEG10(NC-GFP) expression

### CRISPR library generation

For the Perturb-match screen, we created a custom library of 417 sgRNAs targeting 132 genes in known immune signaling pathways, secreted and transmembrane proteins, and signal transduction from extracellular stimuli and 21 safe control intergenic loci (**Supplementary Table 2**). RNA-seq data of LLC cells was used to prioritize genes with TPM>1 *in vitro* at baseline or under IFNγ, IFNβ, or TNF stimulation^6^. Three sgRNAs dropped out in the input pellet and were excluded from analysis. sgRNAs were selected using an ensemble approach, combining scores from CRISPick, Morgens *et al*., VBC, DeepSpCas9, and GuideScan2^89–92^.

### Cancer cell culture

LLC (CRL-1642), 4T1 (CRL-2539), and HEK293T (CRL-3216) cells were purchased from ATCC. LLC, J774, and HEK293T cells were cultured in DMEM (Gibco 11965092) supplemented with 1× GlutaMAX (Gibco 35050061), 100 U mL^−1^ penicillin-streptomycin (Gibco 15140122), and 10% heat-inactivated fetal bovine serum (HI-FBS; Cytiva SH30079.03) and passaged using 0.25% trypsin-EDTA (Gibco 25200114). 4T1 cells were cultured in RPMI 1640 (Gibco 11875093) with the same supplements. All cells were cultured in a humidified 37°C incubator with 5% CO_2_ and normal oxygen and passaged 2-3 times weekly. To generate frozen aliquots, cells were pelleted by centrifugation (250g for 5 min at room temperature), resuspended in bambanker (Bulldog Bio BB05), and frozen at −80°C before transfer to liquid nitrogen for long-term storage. Cell lines were tested and confirmed negative for mycoplasma contamination using MycoAlert (Lonza LT07-218). All cell counting was performed using a Countess 3 automated cell counter (Invitrogen).

### iPSC-derived macrophage differentiation and culture

iPSC-derived macrophages were differentiated from WTC-11 cells (UCSFi001-A, Gladstone Institute, USA). iPSC were cultured in a humidified 37°C incubator with 5% CO_2_ and normal oxygen on Geltrex™ Reduced Growth Factor Basement Membrane Matrix (Gibco A1413302) in mTeSR™-Plus media (STEMCELL Technologies 100-0276) and passaged by single cell suspension after dissociation with Accutase® (Sigma-Aldrich A6964), with supplementation of media with 10 µM Y-27632 (MedChem Express HY-10583) for the first 24h after passaging. All human iPSCs used in this study were previously generated and reported to be derived from material obtained under informed consent and appropriate ethical approvals.

Macrophage differentiation was based upon the previous publications by van Wilgenburg *et al.* and Gutbier *et al.* with several essential modifications^22,93^. Our protocol is as follows: iPSC were recovered from liquid nitrogen storage and passaged at least once before starting differentiation. To initiate mesoderm differentiation, iPSC were lifted with Accutase®, washed once in mTeSR™-Plus or DMEM-F12 (Gibco 11320033) and plated at 1×10^4^ cells per well in a 96-well round bottom ultra-low adhesion plate (Corning 7007) in embryoid body (EB) media (mTeSR™-Plus, 50 ng mL^−1^ BMP4 (Gibco PHC9534), 50 ng mL^−1^ VEGF (Gibco PHC9394), and 20 ng mL^−1^ SCF (R&D 11010-SC-010)) supplemented with 10 µM Y-27632. iPSC were spun at 100g for 3 min with minimal braking during deceleration. Cells were incubated for 48h, before performing a 50% media change with EB media without Y-27632. Cells were then incubated for another 48h and the resulting EBs were then transferred to Factory media (X-VIVO® 15, 1× GlutaMAX, 100 U mL^−1^ penicillin-streptomycin, 50 µM 2-Mercaptoethanol (Gibco 31350010), 25 ng mL^−1^ IL-3 (Peprotech 200-03), and 100 ng mL^−1^ M-CSF (Peprotech 300-25)) in a growth factor reduced Matrigel (Corning 354230) coated (∼6 µg/cm^2^) T175 tissue culture flasks in a final volume of 30 mL and were evenly distributed by gentle shaking. For the first week, 20 mL Factory media was added to the culture every 3-4d, after which a 50% media change was performed every 3-4d until precursors (PreMacs) could be observed in the supernatants. Harvested PreMacs were spun at 400g for 10 min and plated at ∼1×10^5^ cells/cm^2^ in Macrophage media (X-VIVO® 15, 1× GlutaMAX, 100 U mL^−1^ penicillin-streptomycin, and 100 ng mL^−1^ M-CSF) for terminal differentiation into macrophages, with a 50% media change after 3-4d. All cell culture was performed at 37°C, 5% CO_2_, and normal oxygen in a humidified incubator.

### Lentivirus and PEG10 production

For small-scale lentivirus production, 1×10^6^ HEK293T cells were seeded in 6 well plates 18h before transfection. A transfection mixture of 1 µg vector plasmid, 250 ng pMD2.G (Addgene 12259), and 750 ng psPAX2 (Addgene 12260) was prepared in 200 µL of OptiMEM, mixed with 8 µg of PEI pH 7.0 (Polysciences 23966-2), and incubated at room temperature for 20 min. Cells were changed to pre-warmed OptiMEM immediately prior to transfection, and the transfection mixture was added dropwise to cells. Media was changed to complete OptiMEM (5% HI-FBS, 1× MEM non-essential amino acids (Gibco 11140050), 1× GlutaMAX, 50 U mL^−1^ penicillin-streptomycin) 6h after transfection, and the viral supernatant was harvested 24h or 48h afterwards. Cells were transduced at <0.3 multiplicity of infection (MOI).

For large-scale lentivirus production, 36×10^6^ HEK293T cells were seeded in T225 flasks 18h before transfection. A transfection mixture of 42 µg vector plasmid, 13 µg pMD2.G, and 30 µg psPAX2 was prepared in 5 mL of OptiMEM and mixed with 145 µL of Lipofectamine 3000 P3000 reagent. Integrase-deficient lentivirus was generated by substituting psPAX2 with psPAX2(D64V) (Addgene 63586). In a separate tube, 5 mL of OptiMEM was mixed with 165 µL of Lipofectamine L3000 reagent. The two mixtures were combined and incubated at room temperature for 15 min before addition to the HEK293T cells. Media was changed to complete OptiMEM 6h after transfection, and the viral supernatant was harvested 24h or 48h afterwards. All viral supernatants were clarified by centrifugation at 750g for 5 min and filtered through 0.45 µm PVDF filters (Millipore SLHV033RB) before use or storage at −80°C.

PEG10 was generated from a HEK293T PEG10 producer cell line with a stably integrated PEG10 expression cassette under a tetracycline-responsive promoter created by integration of pPD164 using Cp36 large serine recombinase and blasticidin selection for integrants. Producer cells were transfected with pMD2.G and a vector plasmid using Lipofectamine 3000, similar to the large-scale lentivirus protocol above. Cells were changed to complete OptiMEM with 1 µg mL^−1^ doxycycline 6h after transfection, and VLPs were collected and concentrated at 24h and 48h after transfection. All viral supernatants were concentrated using Lenti-X according to the manufacturer’s instructions (Takara Bio 631232) or through ultracentrifugation with G_avg_ of 120,000g for 2h with a sucrose cushion (20% sucrose (w/v), 50 mM Tris-HCl, 100 mM NaCl, 0.5 mM EDTA, pH 7.4) in an Optima XE-90 ultracentrifuge (Beckman-Coulter).

### iPSC-derived macrophage infection and RNA-seq

Lentivirus, integrase-deficient lentivirus, or PEG10 packaging a PAC-T2A-mCherry cargo from pMCB320 was prepared from HEK293T cells using the large-scale production protocol above. iPSCs were differentiated into macrophages and cultured for 1 week with a 50% media change before transduction. For time course experiments, cells were plated in 96-well plates and changed to media with 100 nM SytoxGreen (Invitrogen S7020) and ultracentrifuge-concentrated viral supernatants at various dilutions before imaging every 3h using an Incucyte S3 (Essen).

For bulk RNA-seq experiments, virus dosage was titered to achieve approximately 20% infection rate. Cells were infected in triplicate in 150 mm dishes for 36h before dissociation with Accutase®. The, cells were washed twice with PBS, stained with Live/dead Fixable Aqua (1:1000 dilution, ThermoFisher L34957) for 30 min on ice in the dark, and washed twice with FACS buffer before sorting for mCherry^+^ and mCherry^−^ live cells on a Sony SH800S cell sorter. Total RNA was isolated using the Qiagen RNeasy mini kit according to the manufacturer’s instructions (Qiagen 74104). RNA-seq libraries were sequenced on a NovaSeq 6000 (Illumina). Gene expression was quantified using kallisto pseudoalignment to the hg38 reference genome^94^. Differential gene expression analysis was performed using DESeq2^95^.

### Cell line transfection and transduction

Match-seq sender cell lines were generated by co-transfecting a 1 μg equal mass mixture of pPD430 (Addgene XX) and pAF0084 (Addgene 193462) into 1×10^5^ cells using Lipofectamine 3000 transfection reagent (Invitrogen L3000015) according to the manufacturer’s instructions. Cas9-expressing cells were generated by lentiviral transduction of pPD670. Cells were selected to purity using antibiotic selection for stable integrants.

Pooled CRISPR libraries were lentivirally transduced with 8 μg mL^−1^ polybrene into 3×10^7^ LLC Cas9 sender cells at 7% infection rate, resulting in ∼5,000× coverage per sgRNA. Cells were passaged at 20× unique barcode coverage (100,000× coverage per sgRNA) prior to the tumor challenge and passaged at >20,000× sgRNA coverage for the *in vitro* output arm. An input cell pellet was collected at the time of tumor challenge to record the initial sgRNA distribution, and an *in vitro* output cell pellet was collected at the end of the *in vivo* screen to assess changes in *in vitro* growth caused by perturbations. gDNA from *in vitro* cell pellets was extracted using the Qiagen DNA Blood Midi kit according to the manufacturer’s instructions (Qiagen 51183). The sgRNA-barcode was PCR amplified and sequenced on a MiSeq (Illumina).

Antibiotic selection was performed at the following concentrations: 250 μg mL^−1^ hygromycin B (Gibco 10687010); 10 μg mL^−1^ blasticidin (Gibco A1113903); 10 μg mL^−1^ puromycin (Cayman Chemical 28240). Flow cytometry for fluorescent proteins or HaloTag was used to assess completion of selection (Attune NxT, ThermoFisher).

### *In vivo* tumor challenges

Female C57BL/6J mice ages 6-8 weeks were purchased from Jackson Laboratory. For screen tumor challenges, 4×10^6^ cells were resuspended in 200 μL of a 1:1 mixture of Hanks balanced salt solution (HBSS; Gibco 14025092) and growth factor reduced Matrigel (Corning 356231) and subcutaneously injected into the left flank using a 27G needle. Labeling activity was induced by doxycycline administration *ad libitum* in drinking water at 0.1 mg mL^−1^ for 48h before endpoint tissue harvest. For validation experiments, 2×10^6^ cells were injected in 100 µL of HBSS. For tail vein injections, 500,000 match-seq 4T1 cells with a HaloTag label RNA vector were injected into 6-8 week old female BALB/cJ mice. For cell depletion experiments, mice were injected intraperitoneally with 200 µg anti-CD8β (clone 53-5.8, Bio X Cell), 100 µg anti-CD4 (clone GK1.5, Bio X Cell), 200 µg anti-NK1.1 (clone PK136, Bio X Cell), or 200 µg isotype control (Bio X Cell) every 4d, starting 1d before the tumor challenge. Tumors were measured with calipers every 2-4d, and tumor volume was estimated with the following formula: (length × width^2^ / 2).

All mouse experiments were conducted in accordance with the guidelines of the Institutional Animal Care and Use Committees (IACUC) of Stanford University. For all experiments, no statistical methods were used to predetermine sample sizes. Data collection and analysis were not performed blind to the conditions of the experiment.

### Tumor processing

Tumors for scRNA-seq were dissected and mechanically dissociated using a clean razor blade, followed by enzymatic dissociation for 30 min at 37°C shaking at 250 rpm (1 mg mL^−1^ collagenase IV (Worthington Biochemical NC9919937), 10% dispase (v/v) (Corning 354235), 10% trypsin-EDTA (0.25%) (v/v), and 300 µg mL^−1^ DNase I (Worthington Biochemical LS002006) in HBSS without Ca^++^Mg^++^ (Gibco 14170112)). Tumors for validation experiments were digested using different enzymes to preserve cell surface antigens (1 mg mL^−1^ collagenase A (Roche 10103586001), 2 mg mL^−1^ hyaluronidase (Sigma-Aldrich H3506), and 300 µg mL^−1^ DNase I in RPMI 1640 (Gibco)). Digestions were quenched with ice-cold Leibovitz’s L-15 medium (Gibco 11415064) supplemented with 10% HI-FBS, and cells were passed through a 70 μm strainer. Cells were pelleted at 350g at 4°C and resuspended in ACK lysing buffer (Gibco A1049201) for 1 min. Lysis was quenched using FACS buffer (2% HI-FBS, 4 mM EDTA (Gibco 15575020) in PBS without Ca^++^Mg^++^ (Gibco 10010023)).

Cells were washed twice with PBS with 4 mM EDTA to remove serum, and 1×10^7^ cells from each tumor were resuspended in 170 µL of PBS for MULTI-seq labeling^33^. In brief, a 20 µL mixture of 40 µM equimolar anchor lipid modified oligo (LMO) and barcode oligonucleotides was prepared for each sample. The anchor-barcode mixture was added to the cells from each tumor, briefly vortexed to mix, and incubated on ice for 5 min. Then, 10 µL of 40 µM co-anchor lipid was added to the cells, briefly vortexed, and incubated on ice for 5 min. Labeling was quenched with 1 mL 2% BSA in PBS (w/v) to inactivate unbound LMOs. Samples were pooled and washed twice with 2% BSA.

### Fluorescence-activated cell sorting

Cells were resuspended with TruStain FcX™ PLUS blocking antibody (clone S17011E; BioLegend 156603; 1:100 dilution) and 0.5 µM JF635 HaloTag ligand (ProMega HT1050) in FACS buffer and incubated on ice for 30 min in the dark. Following two washes with FACS buffer, cells were stained using an antibody panel for 30 min in the dark on ice (1:100 dilution unless otherwise noted; anti-CD11b (1:200) (clone M1/70; BD 568345), anti-CD45 (1:500) (clone 30-F11; BioLegend 103116), anti-NK1.1 (clone PK136; BD 741062), Live/dead Fixable Lime (1:1000) (ThermoFisher L34989), anti-Ly76 (clone TER-119; BioLegend 116239), anti-MHC-II (I-A/I-E) (1:500) (clone M5/114.15.2; BioLegend 107643), anti-PDGFRb (clone APB5; BioLegend 136005), anti-CD3 (clone 17A2; BioLegend 100245), anti-CD11c (1:200) (clone N418; BioLegend 117317)). Cells were washed twice with FACS buffer before sorting into chilled 50% HI-FBS in PBS (BD FACSymphony S6). HaloTag-positive cells were gated based on a non-HaloTag-expressing tumor control sample. For scRNA-seq, CD3^+^, NK1.1^+^, CD11b^+^ CD11c^+^ MHC-II^+^, total CD45^+^, and CD45^−^ HaloTag^high^ BFP^+^ cell fractions were sorted in order to control the number of cells from each population loaded for scRNA-seq and enrich for populations that were less abundant in the tumor, such as CD3^+^ and NK1.1^+^ cells (**Extended Data Fig. 3**).

### scRNA-seq library construction

Single-cell suspensions were loaded at ∼100,000 cells per channel into four channels of a Chromium GEM-X 3’ chip (10x Genomics PN-1000690). Transcriptome libraries were prepared according to GEM-X 3’ v4 kit (10x Genomics PN-1000686) instructions using Feature cDNA Primers 2 (10x Genomics PN-2000097) for the cDNA amplification PCR. MULTI-seq libraries were prepared as previously described^33^. All label RNA PCRs were performed using NEB Ultra II Q5 HiFi PCR premix (NEB M0544S) with 30s initial denaturation at 98°C, cycling conditions: denature (98°C, 10s); anneal (variable temperature, 20s); elongation (72°C, 30s), final extension (72°C, 60s), and final hold (4°C). Label RNAs were amplified in PCR1 using 25 µL of cDNA amplification product with 0.5 µM each forward primers oPD910 (TruSeq) and oPD472 (Nextera) and 1 µM biotinylated reverse primer oPD943 (IDT) in 4×50 µL reactions for 12 cycles using 63°C for annealing. PCR products were pooled and cleaned with 0.8× SPRIselect beads (Beckman Coulter B23317) with two 80% ethanol washes, eluting with 30 µL of EB. For each sample, 30 µL of M-280 Dynabeads (ThermoFisher 11205D) were cleaned three times with 1× SSPE buffer (ThermoFisher 15591043) and used to bind PCR1 products for 15 min at room temperature. Bound beads were washed twice with 1× SSPE and once with EB before resuspending in 30 µL of EB^96^. Bead-bound DNA was quantified using a Qubit 3 fluorometer using a High Sensitivity dsDNA kit (ThermoFisher Q32854). PCR2 was performed using all of bead-bound PCR1 products with 0.5 µM each forward primers oPD910 and oPD472 and 0.5 µM reverse primer oPD967 in 2×50 µL reactions for 8 cycles using 67°C for annealing, and the PCR products were cleaned with 1× SPRIselect beads. PCR3 was performed using 10 ng of PCR2 products with 0.5 µM each of forward primers oPD920 and oPD1056 and 0.5 µM of a unique i7 indexing primer in 2×50 µL reactions for 8 cycles using 67°C for annealing, and the PCR products were cleaned with 0.8× SPRIselect beads. PCR amplicon sizes were confirmed using an Agilent 2100 Bioanalyzer using the High Sensitivity DNA kit (Agilent). The resulting library was quantified using qPCR KAPA Library Quantification Kit (Roche 07960140001) and sequenced on a NovaSeq X Plus (Illumina). Primer sequences can be found in **Supplementary Table 17**.

### scRNA-seq data processing

Transcriptome count matrices were generated using Cell Ranger (v9.0.1) by aligning to the mm39 reference transcriptome with Peg10rc4, Cas9-BFP-BSD, VSVG, rtTA-hygroR, and label RNA lentiviral vector synthetic sequences appended. Downstream analysis was performed using Scanpy (v1.10.3)^97^. Cells with >10% mitochondrial gene content or <500 genes detected were removed. Cells were assigned to tumor replicates using MULTI-seq barcodes processed using deMULTIplex2^98^, and droplets with >1 MULTI-seq barcode were called as multiplets and removed. Size factors for each cell *i* were calculated as ∑counts_i_ / mean(∑counts_i_)^99^, and raw counts were transformed by ln(counts / size factor + 1). Principal component analysis (PCA) was performed on transformed gene expression values, and the top 50 PCs were used for nearest-neighbor graph construction and UMAP embedding. Community detection was performed using the Leiden algorithm, and cell type identities were determined based on one-versus-rest differential gene expression. Differential expression analysis between cell clusters was performed using a likelihood ratio test based on logistic regression^100^.

Label RNA reads were processed and error corrected using UMI-tools^101^ (v1.1.2) and aligned to reference libraries (whitelist cell barcodes and sgRNA protospacer sequences) using bowtie (v1.2.2)^102^. Spurious barcodes covered by only one read were discarded, yielding a total of 149,726 sgRNA-randomer combinations. To derive a set of unique barcodes that disallowed barcode sharing between cells from different tumors, a tumor sample identifier was appended to each sgRNA-randomer barcode, resulting in 410,703 unique barcodes. Label RNA matrices were corrected using SCTransform v2 for all downstream analyses^103^.

### match-seq spatial neighborhood inference

To assess the diversity of cell type compositions that were captured by different barcodes, we filtered for barcodes that labeled ≥10 cells and calculated the proportion of each cell type carrying that barcode. A representative subset of 5,000 barcodes was selected by performing geosketch on the barcode × cell type matrix^104^. K-means was used to group barcodes with similar cell type compositions, with K selected by cophenetic coefficient across 30 random initializations. To calculate robust cell type co-localization across replicates, we partitioned cells into spatial groups, calculated the pairwise Pearson correlation coefficient of cell type proportions between spatial groups within each tumor replicate, and performed a two-tailed, one-sample Student’s *t*-test for difference from a null hypothesis of R=0 across the tumor replicates. Spatial groups were calculated by computing the pairwise cosine similarity between cells in the same tumor replicate on the log1p-transformed barcode matrix^105^ and performing community detection using the Leiden algorithm on the fuzzy simplicial graph constructed from the cosine similarity matrix^106,107^.

### sgRNA enrichment analysis

To assign cancer cells to sgRNAs, cells with only a single sgRNA with ≥ 3 UMI counts were assigned to that sgRNA (high confidence). For all remaining unassigned cells (low confidence), we calculated two assignment probabilities: (1) sgRNA identity probability based on a Gamma-Poisson mixed model fitted on sgRNA UMI counts^29^, and (2) a logistic regression model predicting gene perturbation identity based on the top 50 PCs of gene expression fitted on cells with only a single sgRNA with ≥ 5 UMIs. These two probabilities were multiplied by sgRNA UMI counts to generate a ranking statistic. We then assigned sgRNA identity for remaining cells by first checking if the most abundant sgRNA was ≥ *k* UMI counts from the next most abundant sgRNA and then by the previously described ranking statistic to break ties. *k* was set to 5, based on maximizing the marginal increase of the Pearson correlation of sgRNA proportions between the high confidence and the low confidence calls. Enrichment of perturbations was calculated by comparing the number of cancer cells carrying each sgRNA in each tumor to the input pellet sgRNA distribution using MAGeCK RRA (v0.5.9) with 10,000 permutations and normalization to the safe sgRNA distribution^108^.

Significance of label RNA enrichment in receiver cells were calculated using RRA (v0.5.9) by comparing the sgRNA label abundance in each receiver cell type to the sgRNA abundance in the cancer cell population. The mean sgRNA label abundance μ^c^_t_ in each cell type C per tumor T was estimated by fitting a negative binomial distribution across cells. A size factor s^c^_t_ was calculated for each cell type per tumor as mean(μ^c^_t_ / mean(μ^cancer^)), and μ^c^_t_ was divided by s^c^_t_ to size factor normalize. The values were then z-scored using the mean and standard deviation of the safe sgRNA distribution within each tumor, and the difference in z-scores between each receiver cell type and the cancer cells per tumor was taken to generate the statistic r^c^_t_. The mean r^c^_t_ across tumors was taken to generate a ranking statistic r^c^ for each sgRNA. RRA was then used on r^c^ to look for enrichment or depletion of perturbations with 10,000 permutations and normalizing to safe sgRNAs.

### scRNA-seq cancer cell differential gene expression analysis

Differential gene expression of perturbed cancer cells was calculated using a likelihood ratio test based on logistic regression and pooling all cancer cells assigned to an sgRNA targeting a given gene compared to all cells assigned to safe control sgRNA. *P* values were adjusted using the Benjamini-Hochberg procedure^109^, and cNMF^53^ was performed on the signed -log_10_ false discovery rate (FDR), with negative values featurized separately. Features were scaled to unit variance. The consensus solution was selected based on the solution with the lowest distance to K-means centroids of the gene loadings for all random initializations. Pathway enrichment on learned factors was performed by selecting up- and downregulated genes using the kneed package (v0.8.6)^113^ and performing Fisher’s exact test^110,111^.

### scRNA-seq receiver cell differential gene expression analysis

Gamma-Poisson regression models were used to model the effect of perturbed tumor cell interactions on immune cell gene expression^112^. Model coefficients were fitted on pseudobulked cells with tumors with a separate model fitted for each immune cell type with >250 cells. Differential expression compared to total safe sgRNA labels ψ was calculated using contrasts with equal weighting between sgRNAs targeting the same gene. Pathway enrichment scores were calculated using pre-ranked GSEA using the signed F-statistic as the ranking statistic^110,111^. To create gene signatures for interactions by cell type, genes were ranked and top and bottom genes were selected using the kneed package (v0.8.6)^113^. Single cell signature scores were calculated using AUCell for up and down signatures^114^, and the difference between the up and down signature score was used as the final interaction score. The UMAP embedding was calculated on the median value of interaction z-scores across cell states in the following cell types: CD8 T cells, CD4 T cells, NK cells, macrophages, and monocytes.

Differential gene expression matrices for each cell type were factorized by performing cNMF on the signed -log_10_ adjusted *P* values, as described previously. Pathway enrichment on learned factors was performed using pre-ranked GSEA using the factor loadings. To select the NMF factor used for LIA score weight, we selected one factor from each cell type with the highest average enrichment of the following Hallmark gene pathways by signed -log_10_ adjusted *P* value: Inflammatory Response, Interferon Gamma Response, TNFA Signaling via NF-κB, and Allograft Rejection^115^. Perturbation loadings for these factors were scaled to one and averaged for the following cell types to calculate LIA score: CD8 T cells, CD4 T cells, NK cells, macrophages, monocytes, cDC1, cDC2, and MoDC.

### Visium HD 3’ spatial transcriptomics

Tumors were fresh-frozen in Tissue-Tek® optimal cutting temperature compound (OCT) (Sakura Finetek 4583) and stored at −80°C until processing. Tissues were cryosectioned at 10 µm and processed using the Visium HD 3’ kit, according to the manufacturer’s instructions (10x Genomics). Gene expression libraries were sequenced on a NovaSeq 6000 (Illumina). Label RNAs were amplified from the cDNA as described above and sequenced on a NovaSeq X Plus (Illumina). Raw sequencing and image data were processed using Space Ranger (v4.0.1) by aligning to the mm39 reference transcriptome with synthetic sequences appended. Label RNA reads were processed as described above.

### Visium cell type annotation

Cell type composition of the Visium HD 3’ dataset was estimated using cell2location^34^. Reference cell type signatures were inferred from matched dissociated scRNA-seq data using a negative binomial regression model and used for spatial mapping. The following cell types were used as reference classes: cancer, CAF, endothelial, neutrophil, monocyte, MDSC, macrophage, cDC1, cDC2, MoDC, γδ T cell, NK cell, ILC1, Treg, CD8 T cell, CD4 T cell, B cell, pDC, mast cell, basophil, granzyme^high^ cells, and osteoclast-like cells. Bins were mapped to cells using Space Ranger (v4.0.1) single-cell segmentation, with transcript counts aggregated within each segmented cell. The 5th percentile (q05) of the posterior cell type abundance was used as a conservative estimate for each cell.

The following cell types were removed due to low confidence of assignment: γδ T cell, granzyme^high^, ILC1, B cell, pDC, mast cell, and basophil. Dendritic cell annotations cDC1, cDC2 and MoDC were consolidated into a single DC class to obtain 13 cell class annotations. Each cell was assigned to the class with the highest estimated abundance. Cells were retained only when the difference between the highest and second highest abundance exceeded a cell type-specific threshold, with an absolute minimum of 0.05; cells below this margin were excluded from analyses requiring confident assignments. Annotation quality was assessed using curated marker genes and spatial autocorrelation of the corresponding abundance fields.

### Visium barcode spatial analysis

Spatial barcode transfer was quantified separately for each clone, defined by a unique sgRNA-randomer barcode. Cancer cells carrying the barcode were designated source cells, and non-cancer cells carrying the same barcode were designated recipient cells. Because most barcode detections consisted of a single read, barcode-positive cells that did not form a spatially localized cluster were removed using DBSCAN (neighborhood radius, 80 µm; minimum cluster size, three cells) before distance analysis.

For each clone, non-cancer cells were assigned to radial distance bins according to their distance to the nearest source cell. Transfer probability at each distance was calculated as the fraction of cells within each bin carrying the corresponding barcode, thereby accounting for variation in the number of cells available at each distance. Profiles were averaged across the 150 clones with the most recipients, with 95% confidence intervals estimated from 1,000 bootstrap resamples of clones. The distance-dependent transfer probability was fitted as an exponential decay above a constant background in logarithmic space. The resulting half-distance was 46 µm (95% confidence interval, 42-51 µm).

The near-source transfer probability was approximately 6%, corresponding to an approximately 60-fold enrichment over a label permutation background. The estimated half-distance was robust to increasing the detection threshold from one to three reads per cell; higher thresholds increased the enrichment, consistent with single read misassignment attenuation rather than generating the signal. Because the estimated half-distance exceeded the section thickness by several fold and clones extended over millimeter scale distances in the section plane, it represents an upper bound on the true isotropic transfer range.

To characterize the full recipient population, distances from each recipient to the nearest source cell of its clone were pooled across clones and summarized by deciles. This distribution provides a complementary description of the same transfer process and was interpreted together with the transfer probability profile.

Moran’s I for each barcode was calculated on a graph created by Delaunay triangulation on the centroid positions of segmented cells, pruning all edges >50 µm in length. The null distribution was created by permuting reads for each label randomly across all cells for 1000 iterations.

### Tissue staining and microscopy

Tissues were fixed overnight in 4% paraformaldehyde at 4°C, submerged in 30% sucrose overnight, and frozen in OCT at −80°C before cryosectioning onto SuperFrost® Plus slides (Electron Microscopy Sciences 71869-10). Slides were rehydrated by being submerged for 5 min in PBS, 3 min in water, 3 min in 100% ethanol, 3 min in 70% ethanol, and 30s in water. To reduce background autofluorescence, samples were photobleached by immersion in bleaching solution (3% H_2_O_2_, 20mM NaOH in PBS) for 1h at room temperature under a 15,000 lux LED light and washed with PBS with 0.1% Tween 20 (PBST) for 3×5 min. RNA FISH was performed using HCR™ Gold with probes against HaloTag-T2A-PAC following the manufacturer’s instructions (Molecular Instruments). Probe hybridization was performed at 37°C overnight in a humidity chamber. Amplification was performed using X3 with label 647 at room temperature overnight in a humidity chamber. For immunofluorescence staining, samples were blocked in Arnold’s special blocking buffer^116^ (ASBB; 10% HI-FBS, 1% BSA, 0.1% Triton X-100, 0.01% sodium azide in PBS) supplemented with 10% normal donkey serum, for 1h at room temperature before overnight primary antibody staining in ASBB at 4°C in a humidity chamber. The following fluorochrome-conjugated primary antibodies were used: anti-CD45 (1:100 dilution) (clone 30-F11; Biolegend 103144), anti-CD3 (1:50 dilution) (clone 17A2; Biolegend 100212), anti-CD4 (1:50 dilution) (clone RM4-5; Biolegend 100530), anti-CD8α (1:50 dilution) (clone 53-6.7; BD Biosciences 557682), anti-F4/80 (1:50 dilution) (clone D2S9R; Cell Signaling Technology 99651S), anti-ISG15 (1:100 dilution) (clone 1H9L21; Thermo Fisher Scientific 703132RP555), anti-MHC-II (I-A/I-E) (1:100 dilution) (clone M5/114.15.2; Biolegend 107617), anti-ARG1 (1:50 dilution) (clone D4E3M; Cell Signaling Technology 66297S). Samples were washed with PBST for 4×5 min and stained using secondary antibodies (1:500 dilution) and Hoechst 33342 or DAPI (ThermoFisher D1306) in ASBB for 1h at room temperature. Samples were washed with PBST for 3×5 min and mounted with mounting medium (Vector Laboratories H-5501-60) before imaging on a Nikon Eclipse Ti-E spinning disk confocal or a Leica DMi8 inverted epifluorescence microscope. For cyclic immunofluorescence^117^, samples were mounted with 10% glycerol in PBS, and coverslips were demounted after imaging by submerging in PBS for up to 1h. Before subsequent staining rounds, slides were photobleached for 1h at room temperature and washed with PBST for 3×5 min.

### Image processing and quantification

The BaSiC algorithm^118^ was used to calculate dark and flat-field corrections. Images were stitched together and aligned between rounds with ASHLAR^119^. Segmentation masks were generated by running CellPose-SAM^120^ with the default parameters, using the Hoechst stain as the nuclear channel and using either CD3 as the membrane channel for segmenting T cells or F4/80 for segmenting macrophages. All imaging channels across images were independently rescaled by subtracting the 0.5th percentile intensity and dividing by the difference between the 99.5th percentile and the 0.5th percentile (percentile intensities were calculated over all pixels in tumor-containing regions). Single cell measurements were calculated for each cell in the segmentation mask using scikit-image, and these measurements were used to determine whether a cell was positive for a given antibody stain. Specifically, the 80th percentile intensity for each cell was used for antibody stains against plasma membrane proteins (CD3, CD4, CD8α, MHC-II, F4/80), and the mean intensity for each cell was used for antibody stains against cytosolic proteins (ARG1, ISG15). Numeric thresholds to call positive cells were set based on the maximum tumor background intensity, and thresholds were applied uniformly across all samples within the same experiment.

### Quantitative PCR

At least 200,000 sender cancer cells and label-positive immune cells per tumor were sorted using a BD FACSAria III. Cells were immediately lysed using TRI reagent (Zymo Research) and stored in −80°C before processing. RNA was extracted by chloroform extraction and isopropanol precipitation with GlycoBlue co-precipitant (Invitrogen AM9515). Paired DNA was extracted using back-extract buffer (4M guanidine thiocyanate, 1M Tris, 50mM sodium citrate, pH 8.5) and isopropanol precipitation. RNA fractions were digested with ezDNase before reverse transcription using SuperScript IV VILO master mix, according to the manufacturer’s instructions (Invitrogen 11766050). qPCR was performed with triple technical replicates using PrimeTime qPCR probes (IDT) with the following sequences (5’-3’): PAC FAM probe: ACATCGGCAAGGTGTGGGTCG, FWD: TGCAAGAACTCTTCCTCACG, REV: CGATCTCGGCGAACACC; *Ptprc* FAM probe: TGGAGGCTGAATACCAGAGACTTCCT, FWD: TTTCCAATGTGCTGTGTCCT, REV: TGAAGAAGAGAGATCCACCCA; *Rpl13a* probe: ACCTTTGGTCCCCACTTCCCTAGT, FWD: ATGTCCCCTCTACCCACAG, REV: TGAACCCAATAAAGACTGTTTGC; *Ppia* SUN probe: ATGCTGGACCAAACACAAACGG, FWD: TACAGGTCCTGGCATCTTGTCC, REV: CCATTCAGTCTTGGCAGTGCAG. All reactions were performed in a Bio-Rad CFX384 using PrimeTime Gene Expression Master Mix (IDT 1055772). FAM probe Ct values were normalized to the *Ppia* SUN probe and then displayed as fold change over the mean of sender cancer cell samples.

### Creation of CRISPR-edited tumor cell lines for validation studies

Protospacer sequences were cloned into pPD652 and delivered lentivirally into LLC cells with pPD430 and pPD670 cassettes. Integrants were selected using puromycin. Indel formation at the target locus was confirmed by PCR of the target locus from genomic DNA and performing TIDE analysis^121^ and CRISPresso2 analysis^122^ on nanopore sequencing.

Safe sgRNA: GAGTTTGCATCCCTTGTCGT
*Ccm2* sgRNA: TGCTCGGTCCAGAAAGTCGA
*Mif* sgRNA: CACAGCATCGGCAAGATCGG
*Nectin3* sgRNA: ACAATTACTCTGCATAACAT
*Tgfb1* sgRNA: GTACGGCAGTGGCTGAACCA
*Tgfbr1* sgRNA: AGAGCGTTCATGGTTCCGAG
*Traf7* sgRNA: ACCCTGTGATCACTACATGT

## Data availability

*In vivo* Perturb-match data have been deposited in Gene Expression Omnibus under accession code XX. Visium HD 3’ data have been deposited in Gene Expression Omnibus under accession code XX. iPSC-derived macrophage RNA-seq data have been deposited in Gene Expression Omnibus under accession code XX.

## Code availability

Custom code developed for this study is available on GitHub at https://github.com/peterpdu/matchseq_reproduce.

## Acknowledgements

Cell sorting for this project was done on instruments in the Stanford Shared FACS Facility (RRID: SCR_017788) using NIH S10 Shared Instrument Grant (1S10OD026831-01). Sample processing and sequencing of the Visium HD 3’ sample was performed at the Stanford Genomics Core Facility (RRID:SCR_002050). Sequencing was performed at the UCSF Center for Advanced Technology, supported by UCSF PBBR, RRP IMIA, and NIH 1S10OD028511-01 grants. Some of the computing for this project was performed on the Sherlock cluster. We would like to thank Stanford University and the Stanford Research Computing Center for providing computational resources and support that contributed to these research results.

## Funding

This work was supported by funding from the National Cancer Institute (NCI; U54CA261719, U54CA274511), a Lloyd J. Old STAR Award from the Cancer Research Institute, an Emerging Leader Award from the Mark Foundation for Cancer Research, the Stanford Bio-X Interdisciplinary Initiatives Seed Grants Program, and the Parker Institute for Cancer Immunotherapy. A.T.S. is a Weill Cancer Hub West Investigator.

## Author contributions

P.P.D., M.C.B., and A.T.S. conceived and designed the study. P.P.D. cloned all plasmids, created all cell lines, and generated the scRNA-seq dataset. P.P.D. and K.S. cloned the libraries. P.P.D., M.W., and Q.P. analyzed the scRNA-seq and Visium data with input from X.Q. P.P.D., M.P., S.T.K., A.K.M., and M.I.D. performed the mouse studies. S.T.K., P.P.D., and L.H.D. performed the microscopy. A.V.J. prepared the iPSC-derived macrophages. M.C.B., A.T.S., L.B., and P.P.D. acquired funding. P.P.D., M.C.B., and A.T.S. wrote the manuscript, with contributions from all authors. All authors discussed and reviewed the results.

## Competing interests

P.P.D., M.C.B., and A.T.S. are named inventors on a patent related to match-seq. A.T.S. is a founder of Immunai, Cartography Biosciences, Santa Ana Bio, and Arpelos Biosciences, and an advisor to 10x Genomics and Wing Venture Capital.

---

**Extended Data Figure 1.**
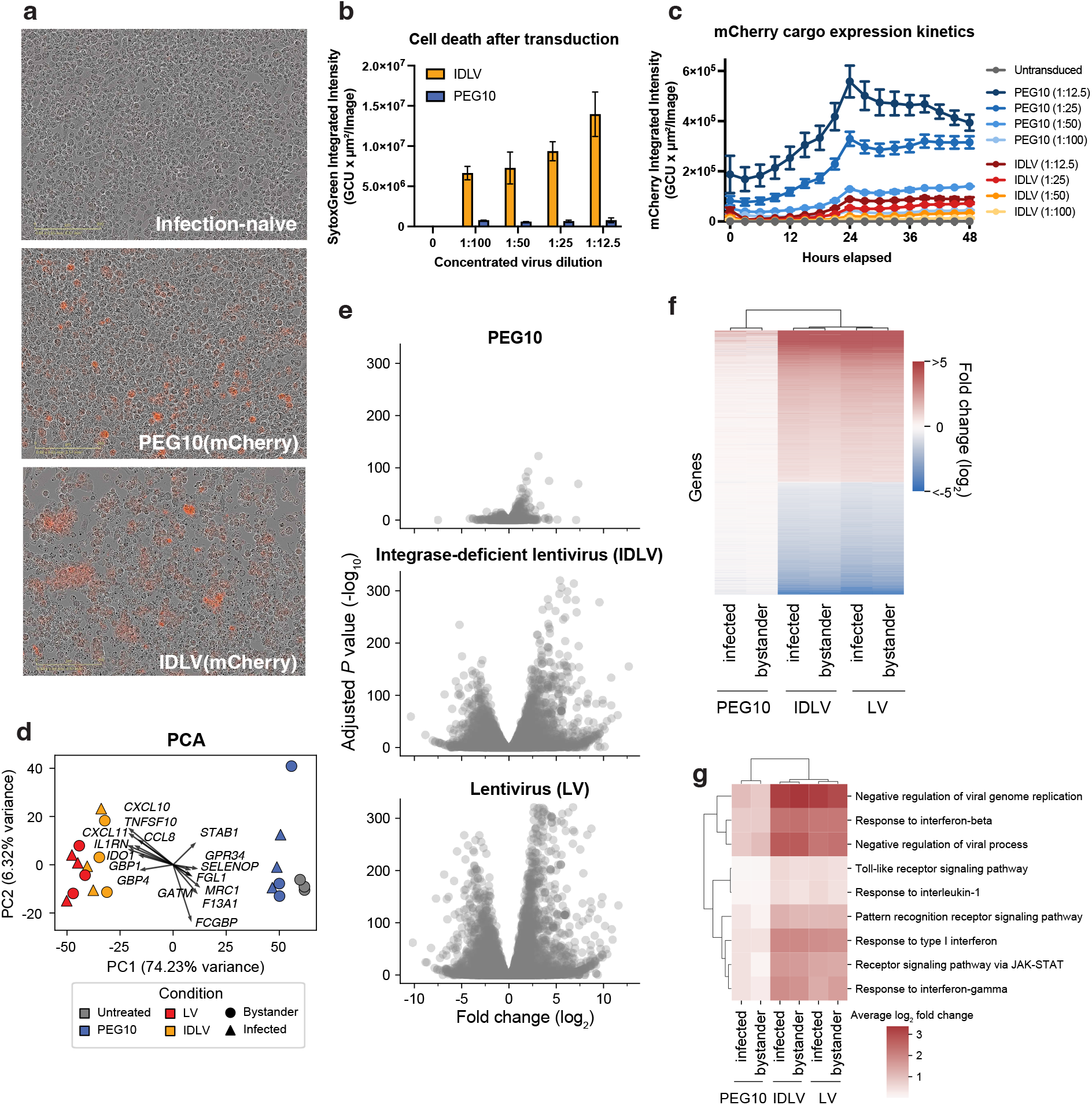
PEG10 efficiently transduces iPSC-derived macrophages with reduced artifactual perturbation compared to lentiviral vectors. A. Representative microscopy images of iPSC-derived macrophages 18h that were either infection-naive (top) or transduced with PEG10 (middle) or IDLV (bottom), showing phase contrast and mCherry fluorescence marking viral delivery. B. Quantification of SytoxGreen integrated intensity marking dead cells of iPSC-derived macrophages treated with a titration of concentrated PEG10 or IDLV packaging an mCherry reporter transcript. Data are means ± s.e.m. of n=3 replicates. C. Time course imaging of mCherry integrated intensity of iPSC-derived macrophages treated with a titration of concentrated PEG10 or IDLV packaging an mCherry reporter transcript. Data are means ± s.e.m. of n=3 replicates. D. PCA biplot of RNA-seq replicate samples. Colors represent infection treatment groups and shapes represent infected (triangle) or bystander (circle) groups. Arrows indicate gene loadings contributing to each principal component. E. Volcano plot showing transcriptome-wide changes comparing iPSC-derived macrophages infected with PEG10 (top), integrase-deficient lentivirus (IDLV, middle), or lentivirus (LV, bottom) after 36h versus infection-naive cells. Values are calculated from n=3 replicates. F. Log_2_ fold change of significantly differentially expressed genes (adj. *P* < 1×10^−10^ and |LFC|>1 in any condition) compared to infection-naive cells. G. Average log_2_ fold change of genes in inflammation and antiviral response gene ontology pathways in PEG10, lentivirus, and IDLV infected cells compared to infection-naive cells.

**Extended Data Figure 2.**
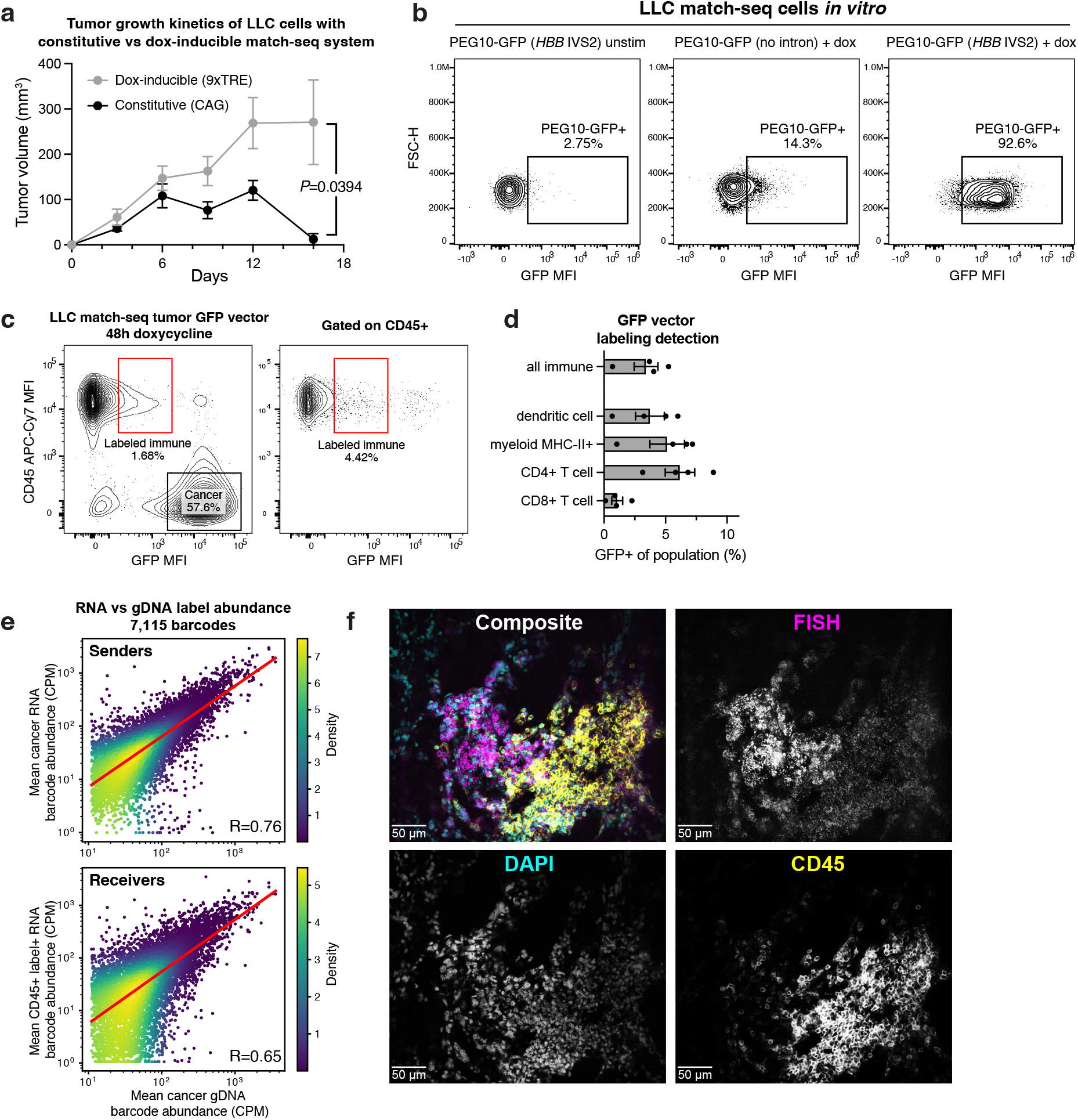
Match-seq optimization and characterization. A. Tumor growth kinetics of LLC cells with constitutively expressed (CAG promoter) or tetracycline-inducible PEG10 and VSVG. Data are means ± s.e.m. n=4 (constitutive) or n=6 (tetracycline-inducible) mice per group. *P* value from Student’s *t*-test. TRE, tetracycline-responsive element. B. *In vitro* inducibility of PEG10-GFP by 48h doxycycline stimulation. Left, unstimulated cells; middle, PEG10 without intron; right, PEG10 with *HBB* IVS2 intron. C. Representative flow cytometry of cells from a LLC match-seq tumor with GFP vector. D. GFP vector labeling detection by flow cytometry of tumor infiltrating immune populations. Data are means ± s.e.m. n=4 mice. E. Barcode sequencing of gDNA fraction of sorted cancer cells (x-axis) and RNA fraction (y-axis) of sorted cancer sender cells (top) or sorted labeled immune cells (bottom). Values are average counts per million across three tumor replicates. Filtered for barcodes that were detected across all replicates and had >10 CPM in the cancer gDNA fraction. F. RNA FISH and immunofluorescent staining of a lung metastasis from 4T1 match-seq cells carrying a HaloTag vector. FISH probes detect the HaloTag-T2A-PAC sequence on the label RNA. CD45^−^ FISH^high^ cells are sender cancer cells and CD45^+^ FISH^low^ cells are labeled immune cells.

**Extended Data Figure 3.**
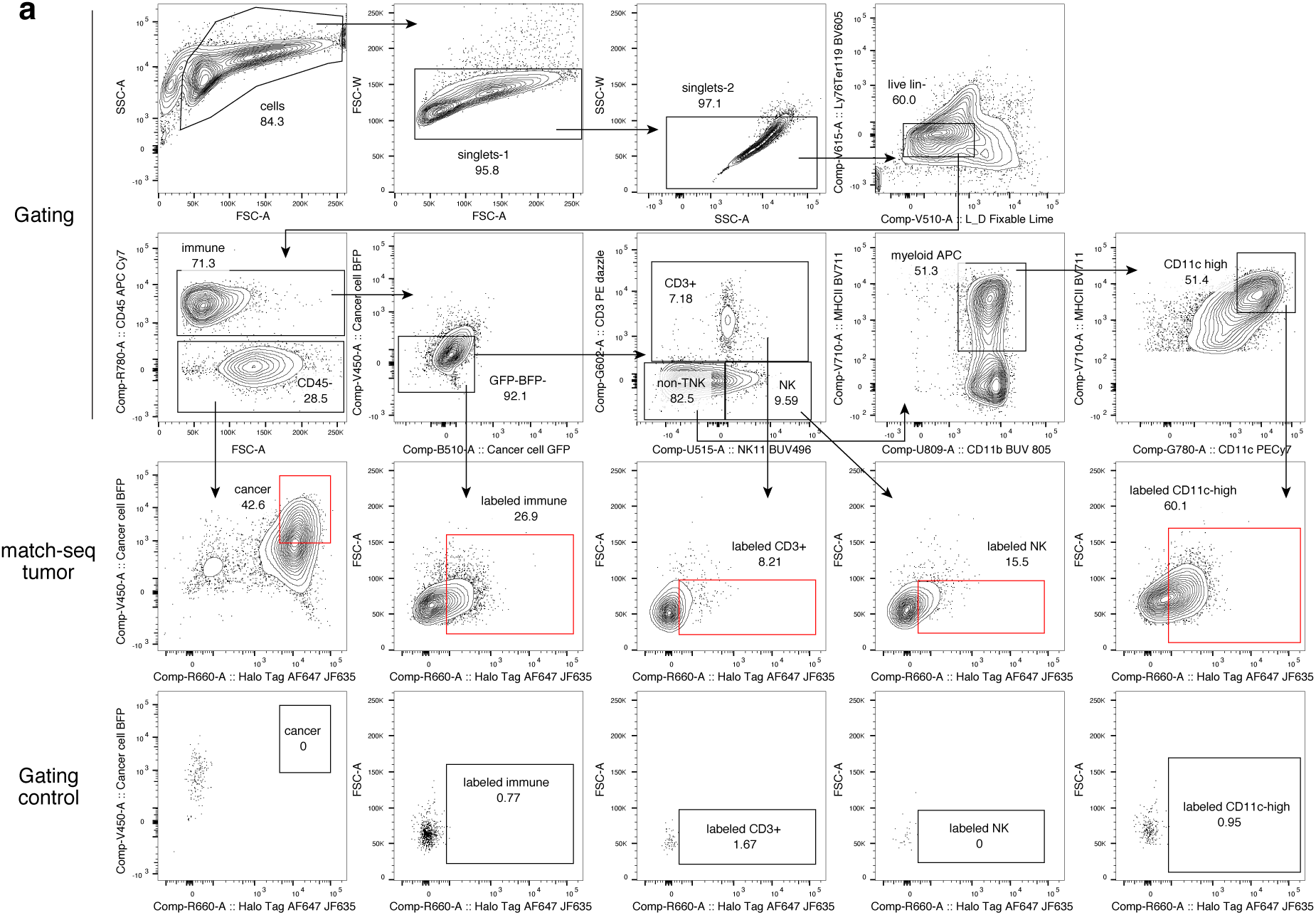
FACS gating scheme for scRNA-seq. A. FACS gating scheme for scRNA-seq to enrich for rare cell populations. Top two rows show cell type gating. The third row shows sort gates for labeled populations and the bottom row shows the gating control for HaloTag signal.

**Extended Data Figure 4.**
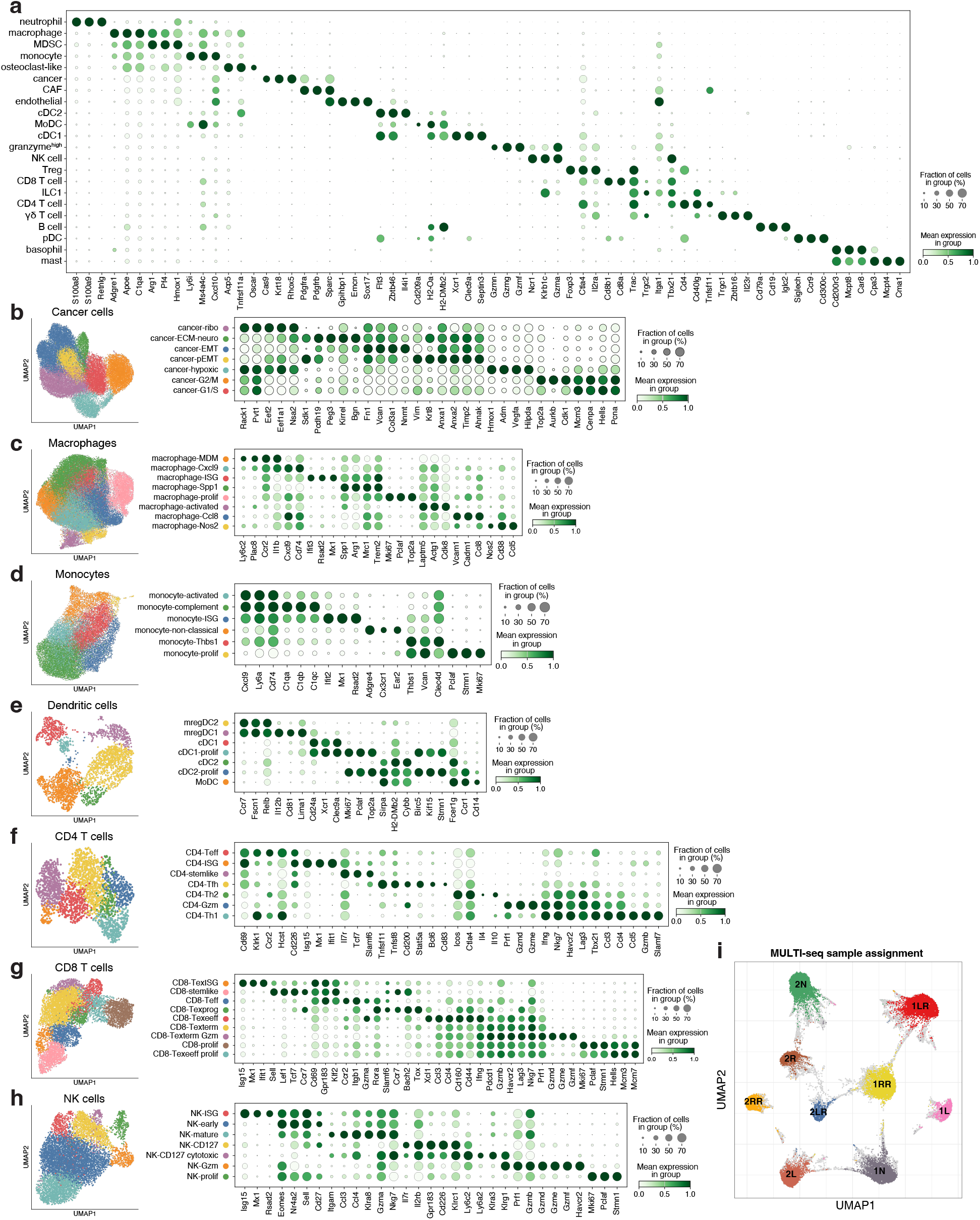
Marker gene expression for cell types and states. A. Marker gene expression for major cell type lineages in the scRNA-seq dataset. Dot size represents the percent of cells expressing the marker gene, and dot color represents the mean expression of the marker gene in the group. CAF: cancer-associated fibroblast; cDC: conventional dendritic cell; ILC1: type 1 innate lymphoid cell; MDSC: myeloid-derived suppressor cell; MoDC: monocyte-derived dendritic cell; NK: natural killer; pDC: plasmacytoid dendritic cell; Treg: regulatory T cell. B. UMAP embedding of cancer cells, colored by cell state (left) and marker gene expression in cell states (right). ECM: extracellular matrix; EMT: epithelial-mesenchymal transition; pEMT: partial epithelial-mesenchymal transition. C. UMAP embedding of macrophages, colored by cell state (left) and marker gene expression in cell states (right). ISG: interferon-stimulated gene; MDM: monocyte-derived macrophage. D. UMAP embedding of monocytes, colored by cell state (left) and marker gene expression in cell states (right). E. UMAP embedding of dendritic cells, colored by cell state (left) and marker gene expression in cell states (right). mregDC: mature dendritic cell enriched in immunoregulatory molecules^87^. F. UMAP embedding of CD4 T cells, colored by cell state (left) and marker gene expression in cell states (right). Teff: effector-like T cell; Tfh: T follicular helper; Th1: T helper type 1; Th2: T helper type 2; Gzm: granzyme. G. UMAP embedding of CD8 T cells, colored by cell state (left) and marker gene expression in cell states (right). Teff: effector-like T cell; Texprog: progenitor exhausted T cell; Texterm: terminally exhausted T cell; Texeeff: early effector exhausted T cell; TexISG: interferon-stimulated gene exhausted T cell. H. UMAP embedding of NK cells, colored by cell state (left) and marker gene expression in cell states (right). I. UMAP embedding of MULTI-seq barcodes with sample barcode assignments.

**Extended Data Figure 5.**
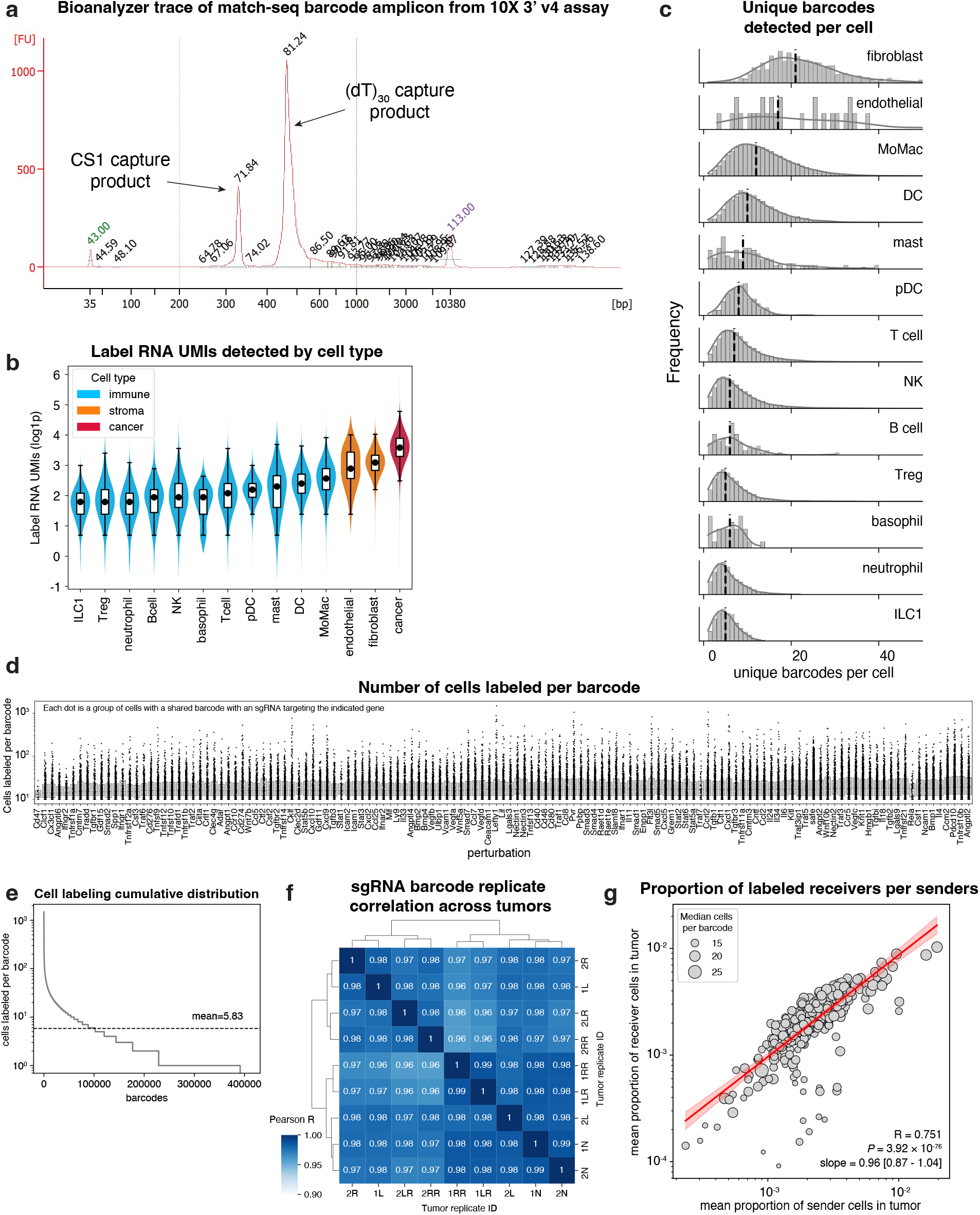
Perturb-match dataset quality metrics. A. Representative bioanalyzer trace for PCR-amplified match-seq barcode amplicon from a scRNA-seq experiment. PCR products from CS1 capture and (dT)_30_ capture are indicated. B. Log1p-transformed UMI counts of label RNAs detected by cell type across the dataset. C. Number of unique barcodes detected by cell type across the dataset. D. Number of cells per unique barcode, grouped by perturbation. Filtered for barcodes with at least 10 cells. E. Cumulative distribution of cells labeled per barcode. Dashed line shows the mean. F. Pairwise Pearson correlation coefficient of log_2_-transformed counts of sgRNA barcodes per tumor. G. Scatter plot showing the correlation between the proportion of sender cells vs the proportion of receiver cells per tumor for an sgRNA. Dot size represents the median number of cells per barcode. Line shows linear fit with 95% confidence interval as the shaded region.

**Extended Data Figure 6.**
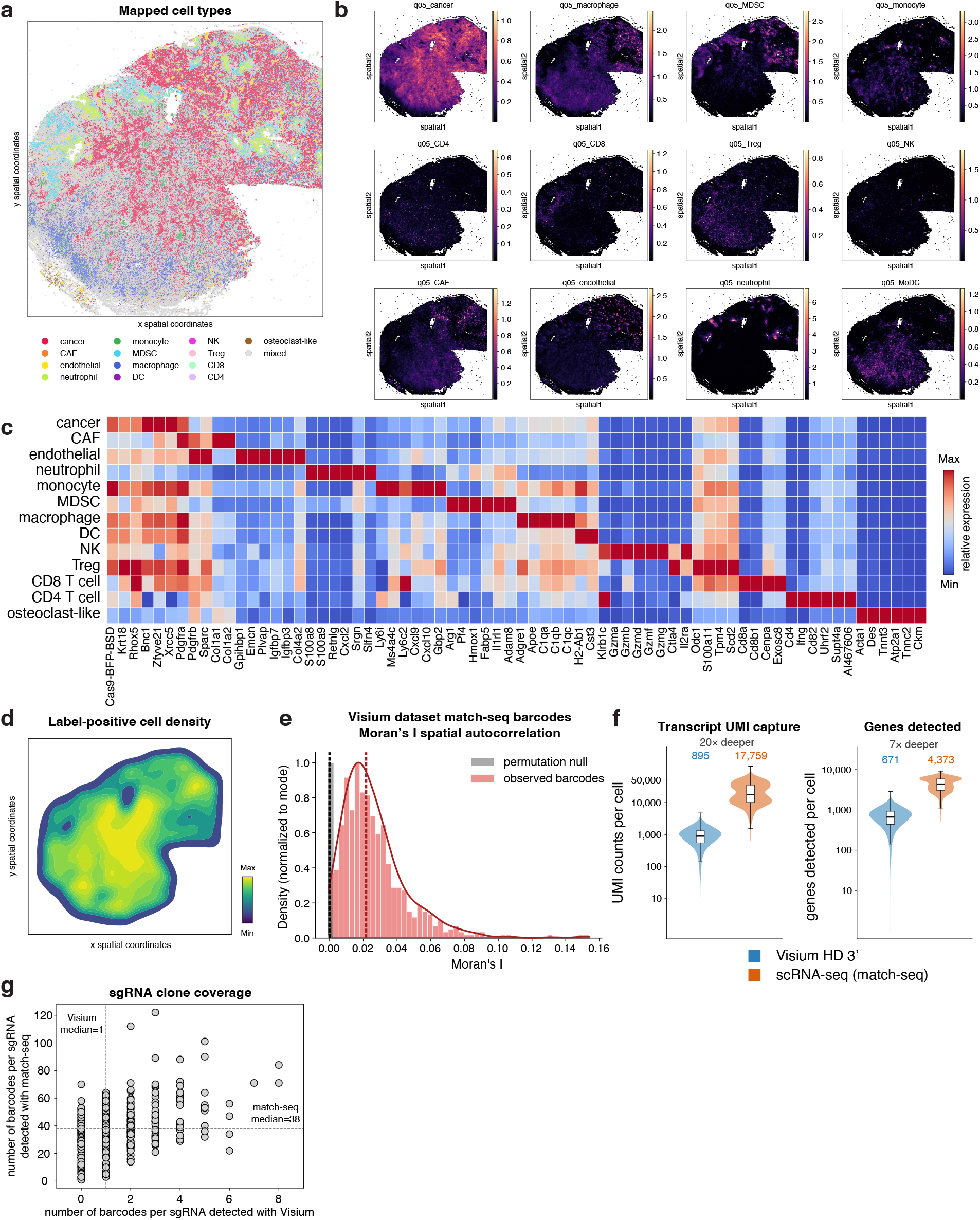
Visium dataset quality metrics. A. Cell types called based on cell2location algorithm. B. The 5th percentile of the posterior distribution from cell2location for reference cell types. C. Marker gene expression of cell types in Visium dataset. D. Kernel density estimate of label-positive cells in spatial coordinates. E. Moran’s I spatial autocorrelation of barcodes in Visium dataset with median indicated by dashed line (red). The grey curve shows spatial permutation null distribution for the mean of 1000 iterations per barcode. The dashed black line shows the Moran’s I expected value. F. Per cell (Left) transcript UMI capture and (right) gene detection in Visium and scRNA-seq datasets for the paired tumor sample. G. Number of barcodes for each sgRNA detected in the Perturb-match dataset or Visium dataset for the paired tumor sample. Dashed lines indicate the median value.

**Extended Data Figure 7.**
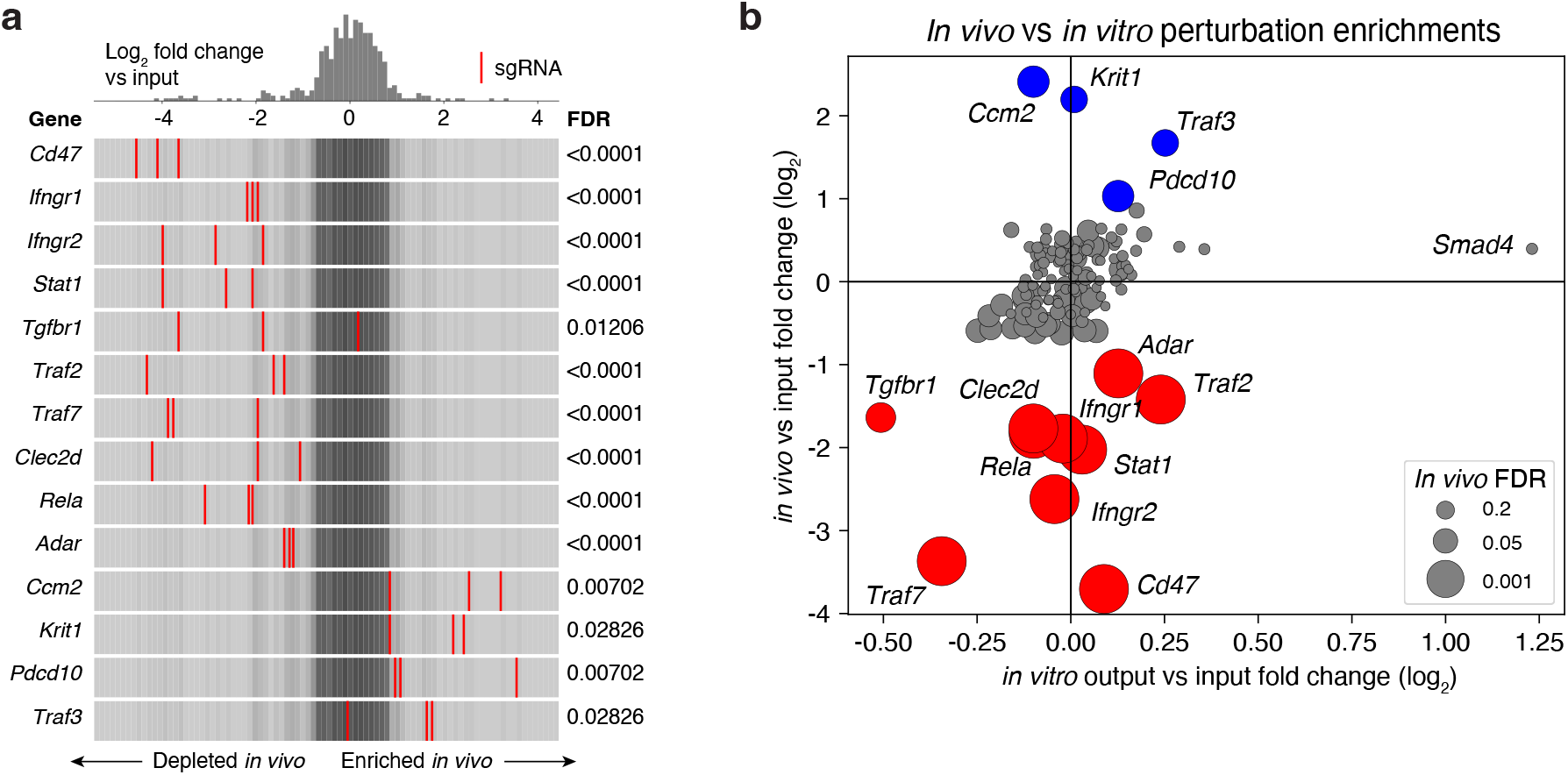
Additional characterization of cancer cell sgRNA enrichment. A. Distributions of sgRNA abundance of cancer cells with the indicated perturbation, normalized to the input sgRNA abundance. FDR from MAGeCK RRA. B. Scatter plot showing log_2_ fold change of mean sgRNA distributions of *in vivo* cells against the *in vitro* output pellet, normalized to input sgRNA abundance. Dot size shows *in vivo* vs input FDR. Blue dots show enriched hits and red dots show depleted hits.

**Extended Data Figure 8.**
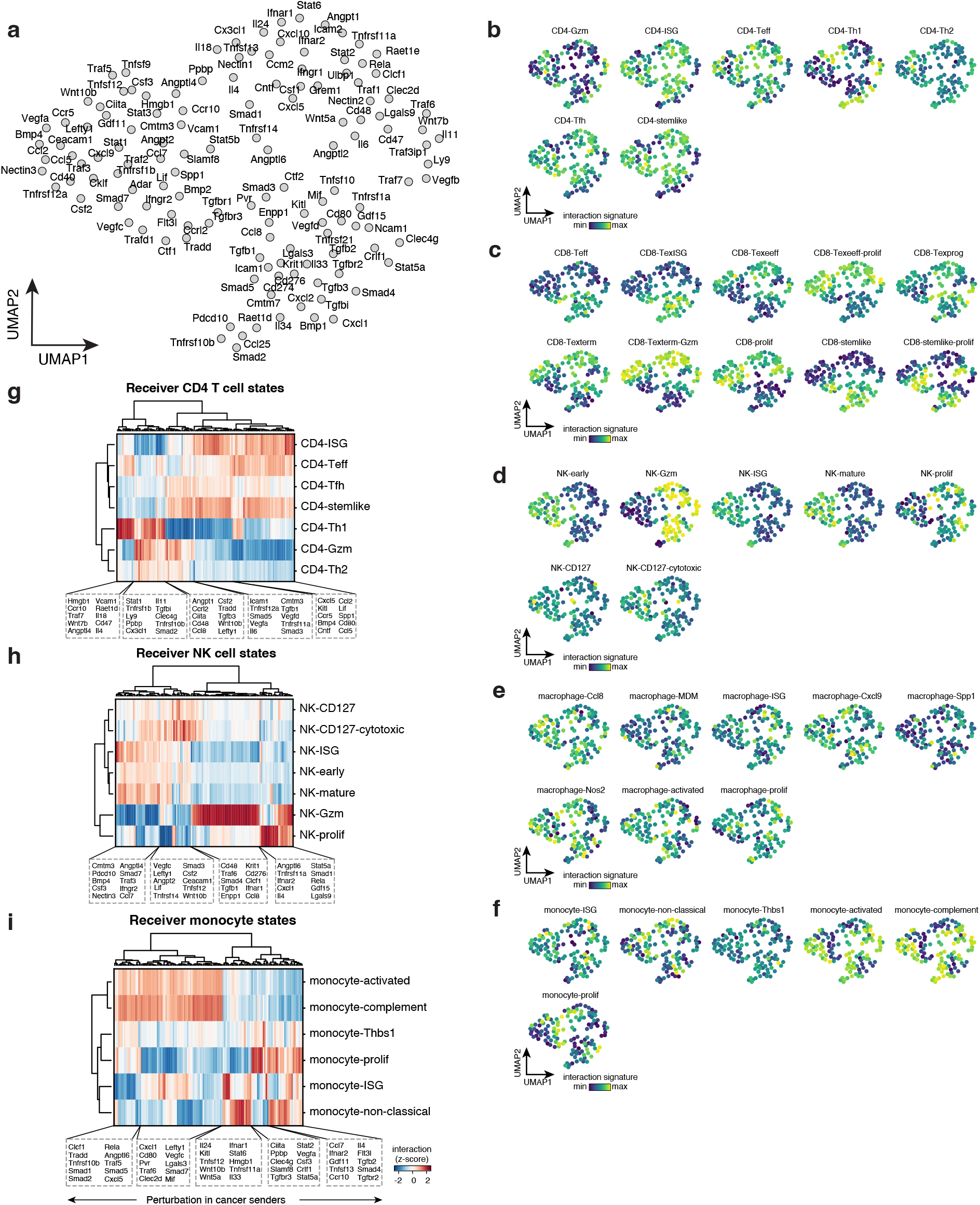
Additional characterization of transcriptional changes in receiver cells upon interaction with perturbed cancer cells. A. UMAP embedding of perturbations based on cell state interaction scores with all points labeled. B. Perturbation UMAPs colored by median cell state interaction scores for all CD4 T cell states. C. Perturbation UMAPs colored by median cell state interaction scores for all CD8 T cell states. D. Perturbation UMAPs colored by median cell state interaction scores for all NK cell states. E. Perturbation UMAPs colored by median cell state interaction scores for all macrophage states. F. Perturbation UMAPs colored by median cell state interaction scores for monocyte states. G. Heatmap showing median interaction scores for cell states in CD4 T cells. H. Heatmap showing median interaction scores for cell states in NK cells. I. Heatmap showing median interaction scores for cell states in monocytes.

**Extended Data Figure 9.**
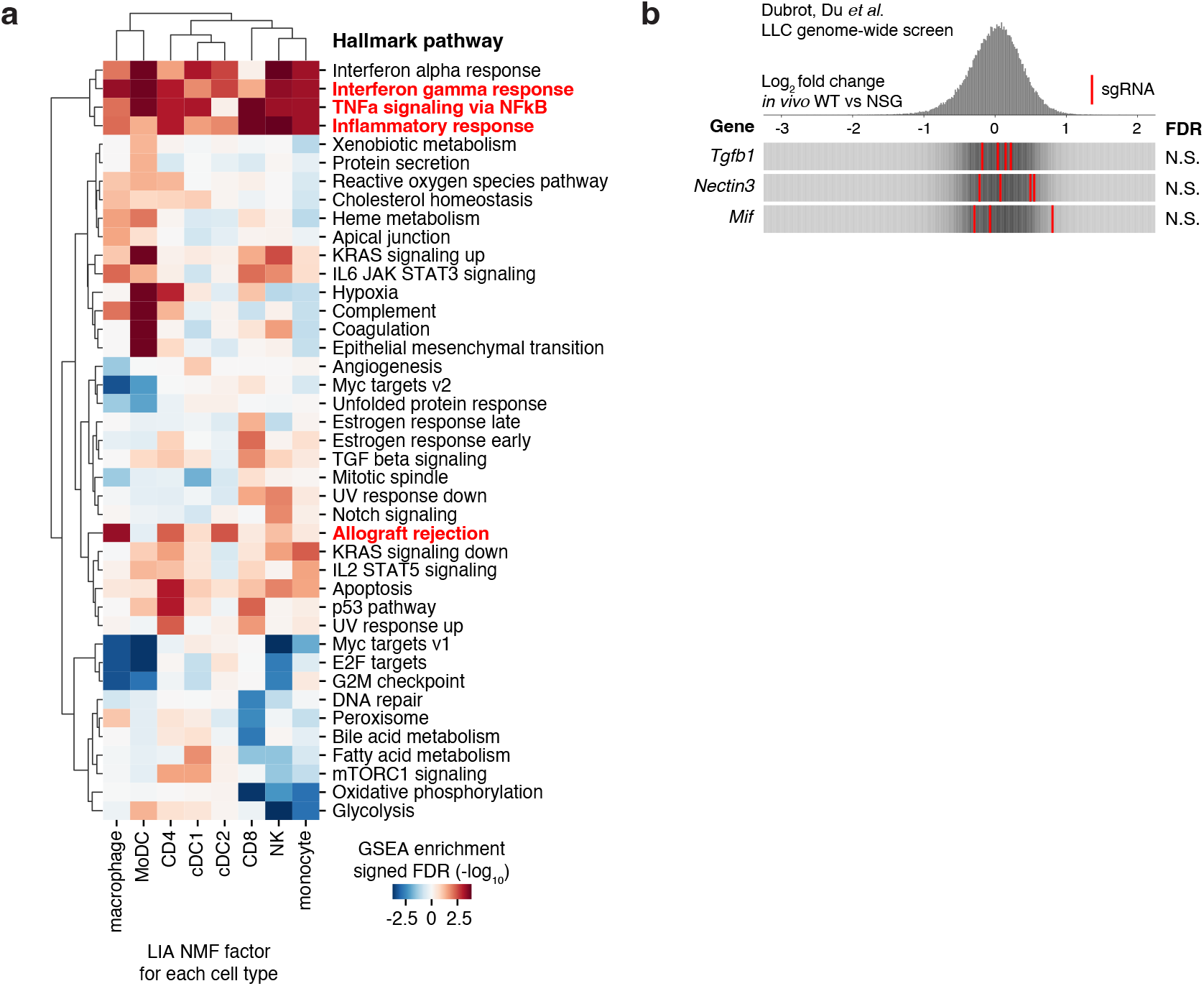
Additional characterization of LIA analysis. A. GSEA pathway enrichment of Hallmark pathways for NMF factors selected from each cell type for LIA score calculation. Gene sets used for assessing local immune activation are highlighted in red. B. Frequency histograms showing log_2_ fold change enrichment or depletion of sgRNA targeting the indicated genes. Data are from Dubrot, Du et al. *in vivo* genome-scale screen in LLC cells comparing immunocompetent WT versus immunodeficient NSG conditions.

## Supplementary Tables

Supplementary Table 1: Differentially expressed genes in iPSC-derived macrophages exposed to lentivirus, integrase-deficient lentivirus, and PEG10

Supplementary Table 2: CRISPR KO library protospacer sequences

Supplementary Table 3: Top 500 marker genes for defining cell states

Supplementary Table 4: Visium barcode labeling statistics

Supplementary Table 5: sgRNA representation in cancer cells in input and *in vivo*

Supplementary Table 6: sgRNA enrichment of cancer cells *in vivo* vs input

Supplementary Table 7: Label representation by sgRNA in receiver cell types

Supplementary Table 8: Label enrichment by sgRNA in receiver cell types

Supplementary Table 9: Cancer cell perturbation differential expression vs safe

Supplementary Table 10: Cancer cell differential expression cNMF loadings values

Supplementary Table 11: Cancer cell cNMF loadings pathway enrichment

Supplementary Table 12: Receiver cell interaction differential expression vs safe

Supplementary Table 13: Receiver cell × perturbation interaction gene signatures

Supplementary Table 14: Receiver cell × perturbation interaction gene signature scores

Supplementary Table 15: Receiver cell interaction cNMF loadings values

Supplementary Table 16: LIA analysis factor loadings

Supplementary Table 17: Oligonucleotide sequences

